# Image processing and analysis methods for the Adolescent Brain Cognitive Development Study

**DOI:** 10.1101/457739

**Authors:** Donald J Hagler, Sean N Hatton, Carolina Makowski, M Daniela Cornejo, Damien A Fair, Anthony Steven Dick, Matthew T Sutherland, BJ Casey, Deanna M Barch, Michael P Harms, Richard Watts, James M Bjork, Hugh P Garavan, Laura Hilmer, Christopher J Pung, Chelsea S Sicat, Joshua Kuperman, Hauke Bartsch, Feng Xue, Mary M Heitzeg, Angela R Laird, Thanh T Trinh, Raul Gonzalez, Susan F Tapert, Michael C Riedel, Lindsay M Squeglia, Luke W Hyde, Monica D Rosenberg, Eric A Earl, Katia D Howlett, Fiona C Baker, Mary Soules, Jazmin Diaz, Octavio Ruiz de Leon, Wesley K Thompson, Michael C Neale, Megan Herting, Elizabeth R Sowell, Ruben P Alvarez, Samuel W Hawes, Mariana Sanchez, Jerzy Bodurka, Florence J Breslin, Amanda Sheffield Morris, Martin P Paulus, W Kyle Simmons, Jonathan R Polimeni, Andre van der Kouwe, Andrew S Nencka, Kevin M Gray, Carlo Pierpaoli, John A Matochik, Antonio Noronha, Will M Aklin, Kevin Conway, Meyer Glantz, Elizabeth Hoffman, Roger Little, Marsha Lopez, Vani Pariyadath, Susan RB Weiss, Dana L Wolff-Hughes, Rebecca DelCarmen-Wiggins, Sarah W Feldstein Ewing, Oscar Miranda-Dominguez, Bonnie J Nagel, Anders J Perrone, Darrick T Sturgeon, Aimee Goldstone, Adolf Pfefferbaum, Kilian M Pohl, Devin Prouty, Kristina Uban, Susan Y Bookheimer, Mirella Dapretto, Adriana Galvan, Kara Bagot, Jay Giedd, M Alejandra Infante, Joanna Jacobus, Kevin Patrick, Paul D Shilling, Rahul Desikan, Yi Li, Leo Sugrue, Marie T Banich, Naomi Friedman, John K Hewitt, Christian Hopfer, Joseph Sakai, Jody Tanabe, Linda B Cottler, Sara Jo Nixon, Linda Chang, Christine Cloak, Thomas Ernst, Gloria Reeves, David N Kennedy, Steve Heeringa, Scott Peltier, John Schulenberg, Chandra Sripada, Robert A Zucker, William G Iacono, Monica Luciana, Finnegan J Calabro, Duncan B Clark, David A Lewis, Beatriz Luna, Claudiu Schirda, Tufikameni Brima, John J Foxe, Edward G Freedman, Daniel W Mruzek, Michael J Mason, Rebekah Huber, Erin McGlade, Andrew Prescot, Perry F Renshaw, Deborah A Yurgelun-Todd, Nicholas A Allgaier, Julie A Dumas, Masha Ivanova, Alexandra Potter, Paul Florsheim, Christine Larson, Krista Lisdahl, Michael E Charness, Bernard Fuemmeler, John M Hettema, Joel Steinberg, Andrey P Anokhin, Paul Glaser, Andrew C Heath, Pamela A Madden, Arielle Baskin-Sommers, R Todd Constable, Steven J Grant, Gayathri J Dowling, Sandra A Brown, Terry L Jernigan, Anders M Dale

## Abstract

The Adolescent Brain Cognitive Development (ABCD) Study is an ongoing, nationwide study of the effects of environmental influences on behavioral and brain development in adolescents. The ABCD Study is a collaborative effort, including a Coordinating Center, 21 data acquisition sites across the United States, and a Data Analysis and Informatics Center (DAIC). The main objective of the study is to recruit and assess over eleven thousand 9-10-year-olds and follow them over the course of 10 years to characterize normative brain and cognitive development, the many factors that influence brain development, and the effects of those factors on mental health and other outcomes. The study employs state-of-the-art multimodal brain imaging, cognitive and clinical assessments, bioassays, and careful assessment of substance use, environment, psychopathological symptoms, and social functioning. The data will provide a resource of unprecedented scale and depth for studying typical and atypical development. Here, we describe the baseline neuroimaging processing and subject-level analysis methods used by the ABCD DAIC in the centralized processing and extraction of neuroanatomical and functional imaging phenotypes. Neuroimaging processing and analyses include modality-specific corrections for distortions and motion, brain segmentation and cortical surface reconstruction derived from structural magnetic resonance imaging (sMRI), analysis of brain microstructure using diffusion MRI (dMRI), task-related analysis of functional MRI (fMRI), and functional connectivity analysis of resting-state fMRI.

**Highlights:**

- An overview of the MRI processing pipeline for the ABCD Study
- A discussion on the challenges of large, multisite population studies
- A methodological reference for users of publicly shared data from the ABCD Study

## Introduction

### ABCD Study Overview

The Adolescent Brain Cognitive Development (ABCD) Study offers an unprecedented opportunity to comprehensively characterize the emergence of pivotal behaviors and predispositions in adolescence that serve as risk or mitigating factors in physical and mental health outcomes (Jernigan et al., 2018; Volkow et al., 2018). Data collection for the ABCD Study was launched in September 2016, with the primary objective of recruiting over 11,000 participants - including more than 800 twin pairs - across the United States over a two-year period. These 9-10-year-olds will be followed over a period of ten years. This age window encompasses a critical developmental period, during which exposure to substances and onset of many mental health disorders co-occur. The ABCD Study is the largest project of its kind to investigate brain development and peri-adolescent health, and includes a comprehensive battery of behavioral assessments (Luciana et al., 2018), multimodal brain imaging (Casey et al., 2018), bioassay data collection (Uban et al., 2018), and other assessments (Bagot et al., 2018; Barch et al., 2018; Lisdahl et al., 2018; Zucker et al., 2018). The longitudinal design of the study, large diverse sample, and open data access policies will allow researchers to address many significant and unanswered questions; for example, understanding the causal interplay between brain development and sleep, exercise, nutrition, and screen time, as well as the contribution of numerous other social, genetic, and environmental factors.

The ABCD consortium is comprised of 21 data acquisition sites in the United States, capturing a nationwide cohort with a diverse and inclusive range of geographic, socioeconomic, ethnic, and health backgrounds. Recruitment of the sample was designed to closely match the demographic profile of the American Community Survey, and the enrollment profiles of 3rd and 4th graders from the National Center for Education Statistics, to achieve a sample reflective of the sociodemographic variation of the US population (Garavan et al., 2018). Two centers oversee the coordination and centralization of resources for the entire consortium, namely the Coordinating Center (CC) and the Data Analysis and Informatics Center (DAIC). Specifically, the CC provides the organizational framework for the scientific and administrative tasks of the ABCD Study and works closely with the DAIC to monitor the progress of each site and the collective consortium towards recruitment targets and other study goals. The DAIC provides the infrastructure for data storage and management and performs centralized processing, curation, and sharing of imaging data. The DAIC is committed to the development of an efficient workflow to lower the barrier to accessing complex image analysis methods for researchers with different skill sets and levels of expertise. The responsibilities of the DAIC include: 1) establishing a harmonized magnetic resonance imaging (MRI) acquisition protocol, with comparable acquisition parameters across scanner vendors; 2) quality control of MRI images before and after processing; 3) centralized image processing and information extraction; 4) public sharing of data and image processing pipelines; and 5) dissemination of imaging-derived measures and tools for use by the consortium and the wider scientific community.

### Large Scale Multimodal Image Acquisition

The past few decades have seen increasing interest in the development and use of noninvasive *in vivo* imaging techniques to study the brain. Rapid progress in MRI methods has allowed researchers to acquire high-resolution anatomical and functional brain images in a reasonable amount of time, which is particularly appealing for pediatric and adolescent populations. The ABCD Study builds upon existing state-of-the-art imaging protocols from the Pediatric Imaging, Neurocognition Genetics (PING) study (Jernigan et al., 2016) and the Human Connectome Project (HCP) (Van Essen et al., 2012) for the collection of multimodal data: T1-weighted (T_1_w) and T_2_-weighted (T_2_w) structural MRI (sMRI), diffusion MRI (dMRI), and functional MRI (fMRI), including both resting-state fMRI (rs-fMRI) and task-fMRI (Casey et al., 2018). The fMRI behavioral tasks include a modified monetary incentive delay task (MID) (Knutson et al., 2000), stop signal task (SST) (Logan, 1994) and emotional n-back task (EN-back) (Cohen et al., 2016). These tasks were selected to probe reward processing, executive control, and working memory, respectively (Casey et al., 2018). The ABCD imaging protocol was designed to extend the benefits of high temporal and spatial resolution of HCP-style imaging (Glasser et al., 2016) to multiple scanner systems and vendors. Through close collaboration with three major MRI system manufacturers (Siemens, General Electric and Philips), the ABCD imaging protocol achieves HCP-style temporal and spatial resolution on all three manufacturers’ 3 Tesla systems without the use of non-commercially available system upgrades.

To help address the challenges of MRI data acquisition with children, real-time motion correction and motion monitoring is used when available to maximize the amount of usable subject data. The prospective motion correction approach, first applied in PING, uses very brief “navigator” images embedded within the sMRI data acquisition, efficient image-based tracking of head position, and compensation for head motion (White et al., 2010). Significant reduction of motion-related image degradation is possible with this method (Brown et al., 2010; Kuperman et al., 2011; Tisdall et al., 2016). Prospective motion correction is currently included in the ABCD imaging protocol for the sMRI acquisitions (T1w and T2w) on Siemens (using navigator-enabled sequences (Tisdall et al., 2012)) and General Electric (GE; using prospective motion (PROMO) sequences (White et al., 2010)) and will soon be implemented on the Philips platform.

Real-time motion monitoring of fMRI acquisitions has been introduced at the Siemens sites using the Frame-wise Integrated Real-time Motion Monitoring (FIRMM) software (Dosenbach et al., 2017). This software assesses head motion in real-time and provides an estimate of the amount of data collected under pre-specified movement thresholds. Operators are provided a display that shows whether our movement criterion (>12.5 minutes of data with framewise displacement (FD (Power et al., 2014)) < 0.2 mm) has been achieved. FIRMM allows the operators to provide additional feedback to participants and also allows operators to adjust their scanning procedures (e.g., skip the final rs-fMRI run) based on whether the criterion has been reached.

### Challenges of Multimodal Image Processing

A variety of challenges accompany efforts to process multimodal imaging data, particularly with large numbers of subjects, multiple sites, and multiple scanner manufacturers. Head motion is a significant issue, particularly with children, as it degrades image quality, and potentially biases derived measures for each modality (Fair et al., 2012; Power et al., 2012; Reuter et al., 2015; Satterthwaite et al., 2012; Van Dijk et al., 2012; Yendiki et al., 2013). As described above, prospective motion correction reduces motion-related image degradation for sMRI, and FIRMM can allow for efficient scanning for fMRI. Despite prospective motion correction for sMRI, image artifacts may persist in children with excessive head motion, so it remains important to include an assessment of motion in data quality reviews. Prospective motion correction for dMRI or fMRI is not yet available for routine use, so post-acquisition volume registration is necessary.

The two main causes of motion-induced artifacts are inconsistencies in the k-space data acquired and violations of the signal model assumptions used in image reconstruction. For example, using an inverse fast Fourier transform image reconstruction assumes the object remained stationary during k-space data sampling. Inconsistencies in k-space acquisition can result in abnormally strong signals or signal loss due to spin dephasing (Zaitsev et al., 2015). For sMRI acquisitions, periodic motion synchronized with the k-space acquisition results in “ghosting” -- a partial or complete replication of the structure along the phase-encoding dimension. Furthermore, motion can introduce blurring and/or ringing, depending on the specifics of the motion, such as its frequency, degree, and location within k-space. In dMRI and fMRI acquisitions, these signal and motion anomalies can result in the subtle corruption of individual frames or slices within a series that can alter derived measures (Liu et al., 2010). Thus, censoring of individual degraded frames and/or slices is one approach used in the literature to minimize contamination of dMRI and fMRI measures (Hagler et al., 2009; Power et al., 2014; Siegel et al., 2014).

The correction of image distortions is another challenge. The single-shot, echo planar imaging (EPI) techniques used for dMRI and fMRI are subject to significant spatial and intensity distortions due to inhomogeneous static magnetic fields, known as B0 distortions. Anatomically accurate, undistorted images are essential for integrating dMRI and fMRI images with anatomical (T_1_w and T_2_w) images, by enabling accurate spatial registration of information across modalities. For longitudinal studies such as the ABCD Study, correcting such B0 distortions is expected to reduce variance in change estimates caused by differences in the precise position of the subject in the scanner. As such, the imaging protocol includes brief spinecho “fieldmap” scans with opposite phase encoding polarities, resulting in opposite spatial distortion patterns, which can be used to correct for B0 distortion. Alignment of the fieldmap images, through nonlinear optimization of deformation fields, enables the removal of spatial and intensity distortions from the EPI images obtained for dMRI and fMRI (Andersson et al., 2003; Chang and Fitzpatrick, 1992; Holland et al., 2010; Morgan et al., 2004).

Another prominent source of spatial distortion in MRI scans is caused by the nonlinearity of the gradient fields used for spatial encoding in MRI (Jovicich et al., 2006; Wald et al., 2001). Distortion due to gradient nonlinearity varies between scanner manufacturers, scanner models, and imaging modes, so distortion correction must be tailored to scanner-type-specific definitions provided by the MRI scanner manufacturers. Such distortions are of particular importance when estimating subtle structural or functional changes from serial MRI scans, since the distortion patterns can vary substantially between scan sessions due to slight differences in subject positioning. Correcting gradient distortions significantly improves the accuracy of longitudinal change estimates based on serial MRI scans (Holland and Dale, 2011).

Eddy current distortion is a significant imaging artifact for dMRI. These distortions, caused by currents induced by diffusion gradients, appear as translation and scaling along the phase-encode direction, with the magnitude and direction of distortions depending on gradient amplitudes and orientations. The resulting misalignment between diffusion-weighted images would, if uncorrected, reduce spatial resolution and cause inaccurate estimation of diffusion metrics (Pierpaoli et al., 1996). There are various implementations of eddy current distortion correction, but a common theme of several approaches is to restrict transformations to displacements in the phase-encode direction (Andersson and Sotiropoulos, 2016; Barnett et al., 2014; Rohde et al., 2004; Zhuang et al., 2006). Eddy current distortion corrections may be further constrained with a spatial transformation model that accounts for diffusion gradient amplitudes and orientations (Zhuang et al., 2006).

An additional challenge of conducting large multi-site longitudinal investigations is addressing image intensity inhomogeneity in structural images, which is particularly problematic when using high density, phased array head coils. The ABCD acquisition sites use either 32 channel head or 64 channel head/neck coils, depending on availability. Standard correction methods, such as those used by FreeSurfer (Dale et al., 1999; Fischl, 2012; Sled et al., 1998) are limited when compensating for steep spatial intensity variation, leading to inaccurate brain segmentation or cortical surface reconstruction. For example, brain tissue farther from the coils, such as the temporal and frontal poles, typically has lower intensity values, causing focal underestimation of the white matter surface, or even resulting in elimination of large pieces of cortex from the cortical surface reconstruction. Furthermore, brain tissue close to coils with extremely high intensity values may be mistaken for non-brain tissue (e.g., scalp). The ABCD image processing pipeline includes an improved intensity inhomogeneity correction of the structural images, using a smoothly varying bias field optimized to standardize image intensities within all white matter voxels (see *sMRI Preprocessing).*

In summary, the ABCD image processing pipeline addresses these known challenges of head motion, distortion and intensity inhomogeneity. Given that the neuroimaging community has been a source of continual development of new methods and improvement and extension of existing methods, we anticipate the emergence over time of improved solutions to these and other challenges. We must also consider future challenges specific to longitudinal analyses, such as the possibility of scanner and head coil upgrade or replacement during the period of study. Therefore, the image processing pipeline is expected to evolve over time to incorporate future improvements and extensions in order to better address the challenges of large-scale multimodal image processing and analysis.

### Data Sharing

The advent of large-scale data sharing efforts and genomics consortia has created exciting opportunities for biomedical research. The availability of datasets drawn from large numbers of subjects enables researchers to address questions not feasible with smaller sample sizes. Public sharing of processing pipelines and tools, in addition to raw and processed data, facilitates replication studies, reproducibility, and meta-analyses with standardized methods, and encourages the application of cross-disciplinary expertise to develop new analytic methods and test new hypotheses. To this end, data and methods sharing is an integral component of the ongoing ABCD Study. Processed data and tabulated region of interest (ROI) based analysis results are made publicly available via the National Institute for Mental Health (NIMH) Data Archive (NDA). Processing pipelines themselves are shared as platform-independent data processing tools.

### Data Management and Curation

Due to the magnitude and complexity of the ABCD Study, data management and curation pose many challenges. Thus, in addition to providing details of the image processing and analysis pipeline, below we also provide brief descriptions of the procedures, processes, and tools critical for managing the flow of the large numbers of imaging datasets from acquisition sites to the DAIC, through the processing pipeline, and into a curated data release. These topics include the transfer of data from sites to the DAIC, MRI protocol compliance checking, visual review of data quality, tracking of missing or corrupted datasets, and the packaging and uploading of processed data to NDA for public release. However, a full description of the design of information systems used for the ABCD Study is beyond the scope of a manuscript focused on the image processing pipeline, and we anticipate that more detailed descriptions of methods related to these topics will be published separately.

## Methods

### Overview

The ABCD DAIC performs centralized processing and analysis of MRI data from each modality, leveraging validated methods used in other large-scale studies, including the Alzheimer’s Disease Neuroimaging Initiative (ADNI) (Mueller et al., 2005), Vietnam Era Twin Study of Aging (VETSA) (Kremen et al., 2010), and PING (Jernigan et al., 2016). We used a collection of processing steps contained within the Multi-Modal Processing Stream (MMPS), a software package developed and maintained in-house at the Center for Multimodal Imaging and Genetics (CMIG) at the University of California, San Diego (UCSD). MMPS provides large-scale, standardized processing and analysis of multimodality neuroimaging data on Linux workstations and compute clusters. MMPS is a toolbox of primarily MATLAB functions, but also includes python, sh, csh scripts, and C++ compiled executables. MMPS also relies upon a number of publicly available neuroimaging software packages, including FreeSurfer (Fischl, 2012), Analysis of Functional NeuroImages (AFNI) (Cox, 1996), and FMRIB Software Library (FSL) (Jenkinson et al., 2012; Smith et al., 2004). The processing pipeline described in this manuscript was used for the ABCD Data Release 1.1, available in October 2018, and a beta testing version has been made publicly available as a self-contained, platform-independent executable (https://www.nitrc.org/projects/abcd_study).

As described above, the ABCD image acquisition protocol includes sMRI, dMRI, and fMRI data. While there are many modality-specific details of the processing and analysis pipeline that will be discussed below, we conceptualize five general stages of processing and analysis (Fig. 1). In the initial unpacking and conversion stage, the DICOM files are sorted by series and classified into types based on metadata extracted from the DICOM headers, and then converted into compressed volume files with one or more frames (time points). DICOM files are also automatically checked for protocol compliance and confirmation that the expected number of files per series was received. In the second stage (processing), images are corrected for distortions and head motion, and cross-modality registrations are performed (Fig. 2). The third stage is brain segmentation, in which the cortical surface is reconstructed and subcortical and white matter regions of the brain are segmented. In the fourth stage (analysis), we carry out modality-specific, single-subject level analyses and extract imaging-derived measures using a variety of regions of interest (ROIs). For the final stage (summarization), ROI analysis results are compiled across subjects and summarized in tabulated form. To provide context for how the centralized image processing is performed in preparation for the public release of data, we will also briefly describe MRI data acquisition, data transfer, quality control, and data and methods sharing. A diagram documenting the various outputs of the processing pipeline can be found in Figure 3.

**Figure 1.**
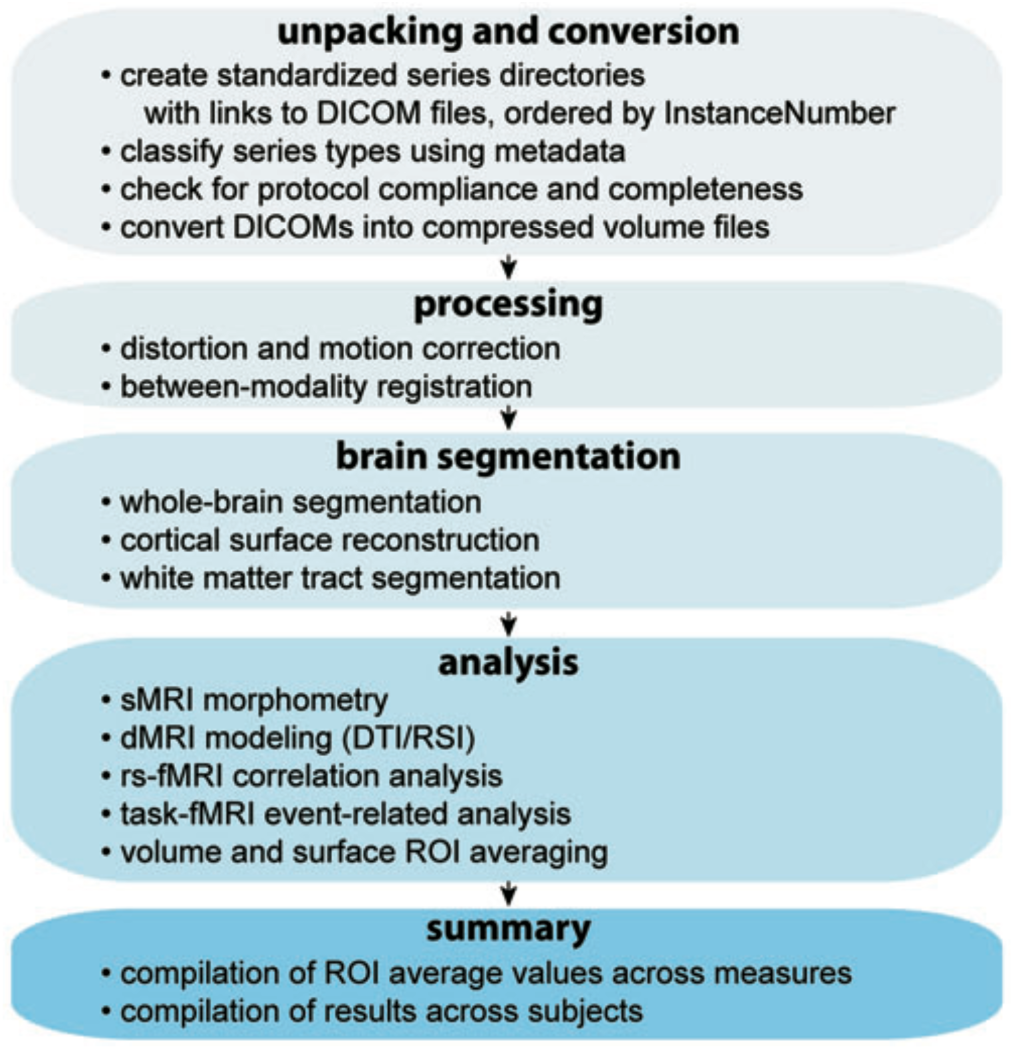
Overview of MMPS processing pipeline steps.

**Figure 2.**
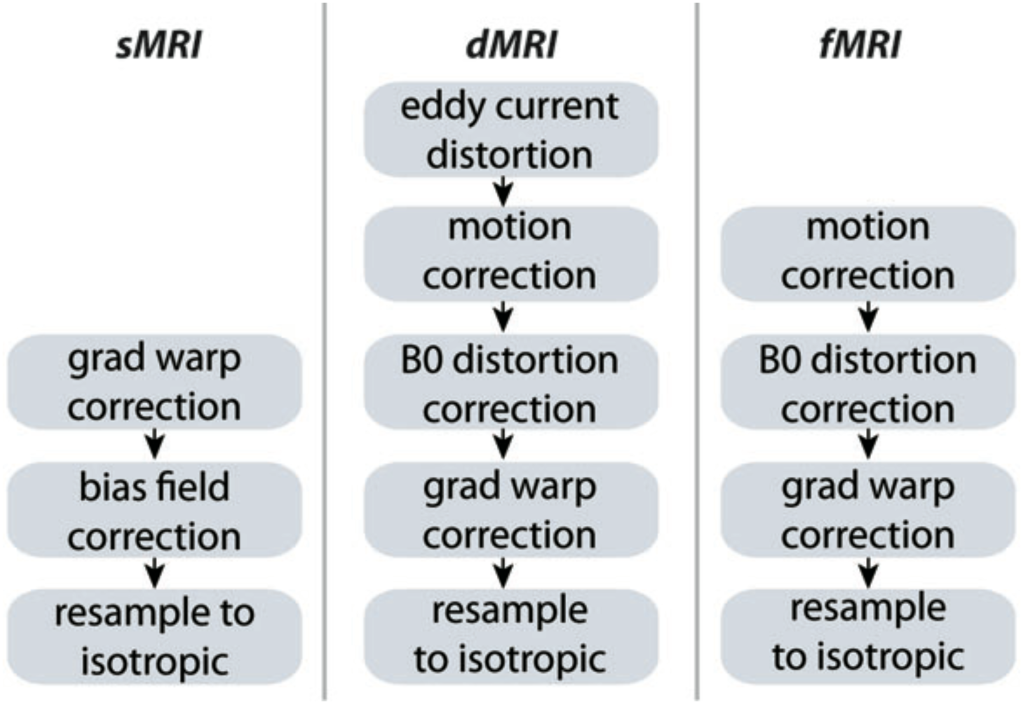
Modality-specific processing steps for bias field, distortion, and/or motion correction.

**Figure 3.**
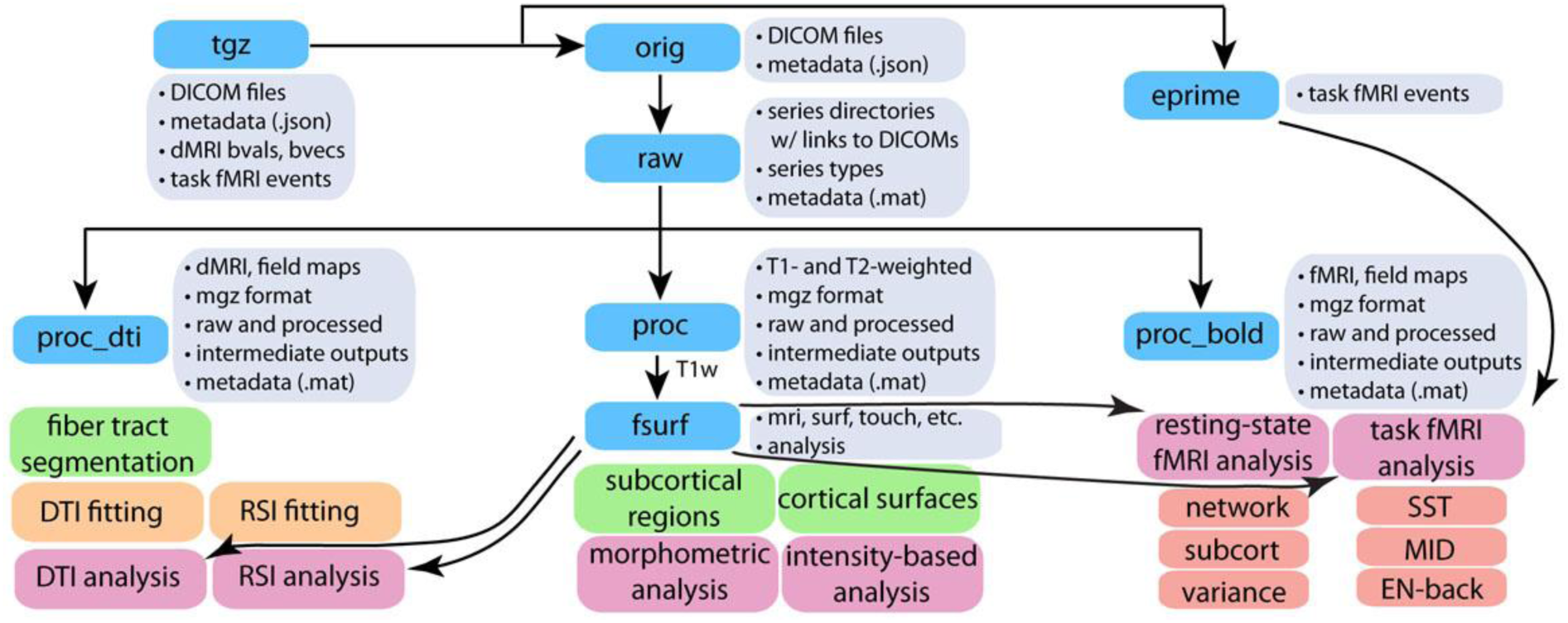
Diagram of MMPS processing pipeline input and outputs.

### Image Acquisition

A standard scan session includes sMRI series (T_1_w and T_2_w), one dMRI series, four rs-fMRI series, and three sets of two task-fMRI series (MID, SST, and EN-back). Minimal details of the imaging protocol are provided here to contextualize the following description of the processing pipeline; additional details have been published previously (Casey et al., 2018). Scan sessions typically require ~2 hours to complete, including a mid-session rest break if the child requests one; they are sometimes split into two separate sessions that take place within one week of each other (3.4% of participants included in ABCD Data Release 1.1). Over 78% of the participants included in ABCD Data Release 1.1 successfully completed the entire image acquisition protocol. Some participants who are unable to complete one or more of the fMRI behavioral tasks (MID, SST, or EN-back), instead perform the missing task or tasks outside the scanner on a laptop computer^1^.

The T_1_w acquisition (1 mm isotropic) is a 3D T_1_w inversion prepared RF-spoiled gradient echo scan using prospective motion correction, when available (Tisdall et al., 2012; White et al., 2010). The T_2_w acquisition (1 mm isotropic) is a 3D T_2_w variable flip angle fast spin echo scan, also using prospective motion correction when available. The dMRI acquisition (1.7 mm isotropic) uses multiband EPI (Moeller et al., 2010; Setsompop et al., 2012) with slice acceleration factor 3 and includes 96 diffusion directions, seven b=0 frames, and four b-values (6 directions with b=500, 15 directions with b=1000, 15 directions with b=2000, and 60 directions with b=3000). The fMRI acquisitions (2.4 mm isotropic, TR=800 ms) also use multiband EPI with slice acceleration factor 6. Each of the dMRI and fMRI acquisition blocks include fieldmap scans for B0 distortion correction. Imaging parameters were made as similar as possible across scanner manufacturers, although some hardware and software constraints were unavoidable (for details, see https://abcdstudy.org/images/Protocol_Imaging_Sequences.pdf).

### Data transfer

Imaging data are transferred from the acquisition sites to the DAIC via dedicated servers, known as FIONA (Flash I/O Network Appliance) Big Data Network Appliances, provided to each scan site by the DAIC. Individual sites push data from their scanner to the local FIONA workstation using the DICOM standard. Data received by the FIONA are automatically checked for completeness and protocol compliance. Protocol compliance checks are performed to ensure that all required scans have been collected and that key parameters such as voxel size or repetition time match expected values for the given scanner. Using a secure web-based application (https://github.com/ABCD-STUDY/FIONASITE, https://scicrunch.org/resolver/SCR_016012) site personnel are able to 1) review data and protocol compliance checks on the FIONA; 2) link the study information to the centralized ABCD database (REDCap, https://www.project-redcap.org) using anonymized identifiers as well as event-related information: and 3) initiate the transfer of data to the DAIC endpoint FIONA device. Data files for each imaging series are packaged separately as compressed archive files (tgz format). Sent along with the imaging data are metadata for each series contained in machine readable text files in JavaScript Object Notation (JSON) format together with md5sum hashes of the tgz files in order to verify the completeness and integrity of data transfer.

After receipt of the imaging data by the DAIC centralized endpoint FIONA device, data are automatically copied to a network attached, high capacity storage device (Synology, Taiwan) using a nightly scheduled rsync operation. The contents of this device are automatically inventoried using the metadata-containing JSON files to categorize series into the different types of imaging series: T-iw, T_2_w, dMRI, rs-fMRI, MID-task-fMRI, SST-task-fMRI, and EN-back-task-fMRI. Missing data are identified by comparing the number of series received of each type to the number of series collected for a given subject, as entered into a REDCap database entry form by the site scan operators. The numbers of received and missing series of each type for each subject are added to the ABCD REDCap database. Using web-based REDCap reports, DAIC staff identify participants with missing data, and either request the data to be re-sent by acquisition sites or address technical problems preventing the data transfer. After receiving imaging data at the DAIC, the tgz files are extracted. For dMRI and fMRI exams collected on the GE platform prior to a software upgrade^2^, this step includes the offline reconstruction of multiband EPI data from raw k-space files into DICOM files, using software supplied by GE.

### sMRI Preprocessing

T_1_w and T_2_w structural images are corrected for gradient nonlinearity distortions using scanner-specific, nonlinear transformations provided by MRI scanner manufacturers (Jovicich et al., 2006). T_2_w images are registered to T_1_w images using mutual information (Wells et al., 1996) after coarse, rigid-body pre-alignment via within-modality registration to atlas brains. MR images are typically degraded by a smooth, spatially varying artifact (receive coil bias) that results in inconsistent intensity variations. Intensity inhomogeneity correction is performed by applying smoothly varying, estimated Brbias fields, using a novel implementation that is similar in purpose to commonly used bias field correction methods (Ashburner and Friston, 2000; Sled et al., 1998). Specifically, B_1_-bias fields are estimated using sparse spatial smoothing and white matter segmentation, with the assumption of uniform T_1_w (or T_2_w) intensity values within white matter. To normalize T_1_w and T_2_w intensities across participants, a target white matter intensity value of 110 is used so that after bias correction, white matter voxel intensities are centered on that target value and all other voxels are scaled relatively. The value of 110 was chosen to match the white matter value assigned by the standard bias correction used by FreeSurfer. The white matter mask, defined using a fast, atlas-based, brain segmentation algorithm, is refined based on a neighborhood filter, in which outliers in intensity -- relative to their neighbors within the mask -- are excluded from the mask. A regularized linear inverse, implemented with an efficient sparse solver, is used to estimate the smoothly varying bias field. The stiffness of the smoothing constraint was optimized to be loose enough to accommodate the extreme variation in intensity that occurs due to proximity to the imaging coils, without overfitting local intensity variations in white matter. The bias field is estimated within a smoothed brain mask that is linearly interpolated to the edge of the volume in both directions along the inferior-superior axis, avoiding discontinuities in intensity between brain and neck.

Images are rigidly registered and resampled into alignment with a custom, in-house atlas brain that has 1.0 mm isotropic voxels and is roughly aligned with the anterior commissure / posterior commissure (AC/PC) axis, facilitating standardized viewing and analysis of brain structure. For most participants, a single scan of each type is collected. If multiple scans of a given type are obtained, only one is used for processing and analysis. Results of manual quality control (QC) performed prior to the full image processing are used to exclude poor quality structural scans (refer to the *Quality Control* section). If there is more than one acceptable scan of a given type, the scan with the fewest issues noted is used. In case of a tie, the final acceptable scan of the session is used.

### dMRI Preprocessing

Eddy current correction (ECC) uses a model-based approach, predicting the pattern of distortions across the entire set of diffusion weighted volumes, based on diffusion gradient orientations and amplitudes, with corrections limited to displacement along the phase-encode direction (Zhuang et al., 2006) as a function of x-, y-, and z-position. With a total of 16 free parameters across the entire set of volumes, we model displacements in the phase encode direction as functions of spatial location, gradient orientation and strength, and frame number (to model a progressive drift observed in some scanners). For each slice of the dMRI volume, a robust tensor fit is calculated in which frames with high residual error -- e.g., signal drop-out in single slices caused by abrupt head motion -- are excluded from the standard linear estimation of tensor model parameters from log transformed images (Basser et al., 1994a). The root mean square (RMS) of the residual error for each frame of each slice is calculated across brain voxels and then normalized by the median RMS value across frames within a given slice. For a given slice, frames with normalized RMS greater than 3.2 are censored from subsequent tensor fits, resulting in a tighter fit for the non-censored frames. A total of three iterations are sufficient to settle upon a stable tensor fit excluding outlier frames for a given slice. To prevent outlier frames from influencing the estimation of eddy current distortions, such frames are replaced (for a given slice) with the corresponding image synthesized from the censored tensor fit. ECC is optimized using Newton’s method through minimization of RMS error between each eddy-current-corrected image and the corresponding image synthesized from the censored tensor fit, accounting for image contrast variation between frames. After applying corrections for the estimated distortions, we re-estimate the tensor, again excluding the outlier frames identified earlier, to produce a more accurate template for subsequent iterations of ECC, with five iterations in total.

To correct images for head motion, we rigid-body-register each frame to the corresponding volume synthesized from the post-ECC censored tensor fit. We remove the influence of outlier frames from motion correction and future analysis by replacing those outlier frame images with values interpolated from the tensor fit calculated without their contribution. The diffusion gradient matrix is adjusted for head rotation, important for accurate model fitting and tractography (Hagler et al., 2009; Leemans and Jones, 2009). Mean head motion values (average FD) are calculated and made available for possible use as covariates in group-level statistical analyses to account for residual effects (Yendiki et al., 2013).

Spatial and intensity distortions caused by B0 field inhomogeneity are minimized using a robust and accurate procedure for reducing spatial and intensity distortions in EPI images that relies on reversing phase-encode polarities (Andersson et al., 2003; Chang and Fitzpatrick, 1992; Holland et al., 2010; Morgan et al., 2004). Pairs of b=0 (i.e., non-diffusion weighted) images with opposite phase encoding polarities (and thus opposite spatial and intensity distortion patterns) are aligned using a fast, nonlinear registration procedure, and the estimated displacement field volume is used to correct distortions in each frame (successive diffusion-weighted volumes) (Holland et al., 2010). Gradient nonlinearity distortions are then corrected for each frame (Jovicich et al., 2006). The b=0 dMRI images are registered to T_1_w structural images using mutual information (Wells et al., 1996) after coarse pre-alignment via within-modality registration to atlas brains. dMRI images are then resampled with 1.7 mm isotropic resolution (equal to the dMRI acquisition resolution), with a fixed rotation and translation relative to the corresponding T_1_w image that has been rigidly resampled into alignment with an atlas brain. This provides a standard orientation for the resulting dMRI images, fitting the brain within the set of axial dMRI slices and producing more consistent diffusion orientations across participants, as viewed with diffusion encoded color (DEC) fractional anisotropy (FA) map images. The diffusion gradient matrix is again adjusted for head rotation. Cubic interpolation is used for each of these resampling steps. A registration matrix is provided to specify the rigid-body transformation between dMRI and T_1_w images.

### fMRI Preprocessing

Head motion is corrected by registering each frame to the first using AFNI’s 3dvolreg (Cox, 1996), which also provides estimates of head motion time courses that are incorporated into task-fMRI and resting-state-fMRI single-subject analyses (see below). B_0_ distortions are corrected using the same reversing polarity method used for the dMRI (Holland et al., 2010). To avoid signal “drop-out” due to within-voxel field gradients in gradient-echo acquisitions, the displacement field is estimated from separate spin-echo calibration scans, then adjusted for estimated between-scan head motion, and finally applied to the series of gradient-echo images. Images are next corrected for distortions due to gradient nonlinearities (Jovicich et al., 2006). Finally, all fMRI scans for a given participant’s imaging visit are resampled with cubic interpolation into alignment with each other, correcting between-scan motion, using a scan in the middle of the session as the reference. Automated registration between the spin-echo, B0 calibration scans (i.e., field maps) and T_1_w structural images is performed using mutual information (Wells et al., 1996) with coarse pre-alignment based on within-modality registration to atlas brains. A registration matrix is provided to specify the rigid-body transformation between fMRI and T_1_w images. The resulting fMRI images remain in “native-space” and have 2.4 mm isotropic resolution.

### Brain Segmentation

Cortical surface reconstruction and subcortical segmentation are performed using FreeSurfer v5.3.0, which includes tools for estimation of various measures of brain morphometry and uses routinely acquired T_1_w MRI volumes (Dale et al., 1999; Dale and Sereno, 1993; Fischl and Dale, 2000; Fischl et al., 2001; Fischl et al., 2002; Fischl et al., 1999a; Fischl et al., 1999b; Fischl et al., 2004; Segonne et al., 2004; Segonne et al., 2007). The FreeSurfer package has been validated for use in children (Ghosh et al., 2010) and used successfully in large pediatric studies (Jernigan et al., 2016; Levman et al., 2017). Cortical surface reconstruction includes skull-stripping (Segonne et al., 2004), white matter segmentation, initial mesh creation (Dale et al., 1999), correction of topological defects (Fischl et al., 2001; Segonne et al., 2007), generation of optimal white and pial surfaces (Dale et al., 1999; Dale and Sereno, 1993; Fischl and Dale, 2000), and nonlinear registration to a spherical surface-based atlas based on the alignment of sulcal/gyral patterns (Fischl et al., 1999b). Because intensity scaling and inhomogeneity correction are previously applied (refer *sMRI Preprocessing),* the standard FreeSurfer pipeline was modified to bypass the initial intensity scaling and N3 intensity inhomogeneity correction (Sled et al., 1998). As of ABCD Data Release 1.1, the T_2_w MRI volumes are not used in the cortical surface reconstruction and subcortical segmentation, but this may be incorporated in future releases.

Subcortical structures are labeled using an automated, atlas-based, volumetric segmentation procedure (Fischl et al., 2002) (Supp. Table 1). Labels for cortical gray matter and underlying white matter voxels are assigned based on surface-based nonlinear registration to the atlas based on cortical folding patterns (Fischl et al., 1999b) and Bayesian classification rules (Desikan et al., 2006; Fischl et al., 2004) (Supp. Table 2). Fuzzy-cluster parcellations based on genetic correlation of surface area are used to calculate averages of cortical surface measures for each parcel (Chen et al., 2012) (Supp. Table 3). Functionally-defined parcels, based on resting-state correlations in fMRI (Gordon et al., 2014), are resampled from atlas-space to individual subject-space, and used for resting-state fMRI analysis (Supp. Table 4).

Major white matter tracts are labelled using AtlasTrack, a probabilistic atlas-based method for automated segmentation of white matter fiber tracts (Hagler et al., 2009). The fiber atlas contains prior probabilities and orientation information for specific long-range projection fibers, including some additional fiber tracts not included in the original description (Hagler et al., 2009), such as cortico-striate connections and inferior to superior frontal cortico-cortical connections (Supp. Table 5). sMRI images for each subject are nonlinearly registered to the atlas using discrete cosine transforms (DCT) (Friston et al., 1995), and diffusion tensor imaging (DTI)-derived diffusion orientations for each subject are compared to the atlas fiber orientations, refining the *a priori* tract location probabilities, individualizing the fiber tract ROIs, and minimizing the contribution from regions inconsistent with the atlas. Voxels containing primarily gray matter or cerebral spinal fluid, identified using FreeSurfer’s automated brain segmentation (Fischl et al., 2002), are excluded from analysis.

### sMRI Morphometric and Image Intensity Analysis

Morphometric measures include cortical thickness (Fischl and Dale, 2000; Rimol et al., 2010), area (Chen et al., 2012; Joyner et al., 2009), volume, and sulcal depth (Fischl et al., 1999a). Image intensity measures include T_1_w, T_2_w, and T_1_w and T_2_w cortical contrast (normalized difference between gray and white matter intensity values) (Westlye et al., 2009). We sample intensity values at a distance of ±0.2 mm -- relative to the gray-white boundary-along the normal vector at each surface location and calculate cortical contrast from gray and white matter values ([white - gray] / [white + gray] / 2). We calculate averages for each cortical parcel in the default FreeSurfer parcellation scheme (Desikan et al., 2006) using unsmoothed, surface-based maps of morphometric and image intensity measures. For each of the fuzzy-cluster parcels (Chen et al., 2012), we calculate weighted averages (weighted by fuzzy cluster membership values ranging from 0 to 1) for each measure using smoothed surface maps (~66 mm FWHM, matching the level of smoothing used for derivation of the fuzzy cluster parcels). We also calculate averages of the unsmoothed intensity measures for the volumetric subcortical ROIs, in addition to the volume of each structure.

### dMRl Microstructural Analysis

We calculate several standard measures related to microstructural tissue properties using DTI (Basser et al., 1994b; Basser and Pierpaoli, 1996), including FA and mean, longitudinal (or axial), and transverse (or radial) diffusivity (MD, LD, and TD). Diffusion tensor parameters are calculated using a standard, linear estimation approach with log-transformed diffusion-weighted (DW) signals (Basser et al., 1994a). Frames with b>1000 are excluded from tensor fitting (leaving 6 directions with b=500, 15 directions with b=1000) so that the DTI-derived measures better correspond to those derived from traditional, single-b-value acquisitions (and better correspond to the assumptions of the DTI model). Tensor matrices are diagonalized using singular value decomposition, obtaining three eigenvectors and three corresponding eigenvalues. FA is calculated from the eigenvalues, as described elsewhere (Basser and Pierpaoli, 1996). MD is calculated as the mean of the eigenvalues. LD is the first eigenvalue, and TD is the mean of the second and third eigenvalues (Alexander et al., 2007).

Taking advantage of the multiple b-value acquisition, we also fit a Restriction Spectrum Imaging (RSI) model (White et al., 2013a; White et al., 2014; White et al., 2013b), a linear estimation approach that allows for mixtures of “restricted” and “hindered” diffusion within individual voxels. We use RSI to model two volume fractions, representing intracellular (restricted) and extracellular (hindered) diffusion, with separate fiber orientation density (FOD) functions, modeled as fourth order spherical harmonic functions, allowing for multiple diffusion orientations within a single voxel. For both fractions, LD is held constant, with a value of 1 × 10^-3^ mm^2^/s. For the restricted fraction, TD is modelled as 0. For the hindered fraction, TD is modelled as 0.9 × 10^−3^ mm^2^/s. Measures derived from this RSI model fit (summarized in Supp. Table 6) include the following: restricted normalized isotropic (N0), restricted normalized directional (ND), restricted normalized total (NT), hindered normalized isotropic (N0_s2), hindered normalized directional (ND_s2), and hindered normalized total (NT_s2). Each of these measures is defined as the Euclidean norm (square root of the sum of squares) of the corresponding model coefficients divided by the norm of all model coefficients. These normalized RSI measures are unitless and range from 0 to 1. The square of each of these measures is equivalent to the signal fraction for their respective model components. N0 and NT_s2 are derived from the 0th order spherical harmonic coefficients of the restricted and hindered fractions, respectively, and reflect varying contributions of intracellular and extracellular spaces to isotropic diffusion-related signal decreases in a given voxel. ND and ND_s2 are calculated from norm of the 2nd and 4th order spherical harmonic coefficients of the restricted and hindered fractions, respectively. These higher order components reflect oriented diffusion; diffusion that is greater in one orientation than others. Qualitatively, ND is very similar to FA, except that, by design, it is unaffected by crossing fibers. NT and NT_s2 reflect the overall contribution to diffusion signals of intracellular and extracellular spaces, and are calculated from the norm of the 0th, 2nd, and 4th order coefficients of the restricted and hindered fractions, respectively, again divided by the norm of all model coefficients.

Mean DTI and RSI measures are calculated for white matter fiber tract ROIs created with AtlasTrack and for ROIs derived from FreeSurfer’s automated subcortical segmentation. With fiber tracts represented as thresholded probability maps, probability estimates are used to calculate weighted averages of DTI and RSI measures. DTI and RSI measures are also sampled onto the FreeSurfer-derived cortical surface mesh to make maps of diffusion properties for cortical gray matter and white matter adjacent to the cortex (Govindan et al., 2013; Kang et al., 2012) and calculate surface-based ROI averages. Values are sampled with linear interpolation perpendicular to the gray/white boundary (“white” surface) in 0.2 mm increments, ranging from 0.8-2 mm in both directions. White and gray matter values are calculated by combining samples within tissue type using a weighted average based on the proportion of white or gray matter in each voxel (Elman et al., 2017). For subcortical ROIs, contamination due to partial voluming in the ROI with CSF is suppressed by calculating weighted averages. Specifically, weighting factors for each voxel in the ROI are calculated based on the difference of MD values relative to the median within each ROI. The typical dispersion of MD values is defined for each ROI as the median absolute deviation from the median (MAD), averaged across subjects. Weighting factors are calculated using Tukey’s bisquare function such that lower weights are assigned to voxels with MD values farther from the median value, relative to the dispersion values multiplied by 4.7 (Tukey, 1960).

### Task-fMRI Behavioral Measures

Behavioral measures specific to each of the three tasks are calculated to assess the performance of the participant during the task and identify participants with poor accuracy or slow reaction times (Casey et al., 2018). For the MID task, there are three types of trials; participants have a chance to either win money, lose money, or earn nothing. Wins and losses are further subdivided into small and large magnitudes. After a short response time window, positive or negative feedback informs the participant about performance in each trial. The behavioral metrics for each type of MID trial are: the number of trials, mean and standard deviation (SD) of the reaction times to the different incentive magnitudes, and total monetary earning. For the SST, participants are asked to indicate by button press if the direction of an arrow presented is leftward or rightward during a short response time window; this is a Go trial. On some trials, a signal is subsequently presented to withhold the motor response, making it a Stop trial. The primary categories of trial-response combinations are “Correct Go”, “Incorrect Go”, “Correct Stop”, and “Incorrect Stop”. Additional, typically rare, categories are “Correct Late Go” (a “Late Go” response is defined as >1000 ms after arrow presentation), “Incorrect Late Go”, “Stop Signal Delay” (response during interval between arrow presentation and stop signal onset), and “No Response”. For each category, number of trials and mean and SD of the reaction times are provided. Additional metrics include mean stop signal delay and mean stop signal reaction time. The EN-Back task is a block design of 0-back and 2-back working memory tasks in which participants are asked to respond to emotionally positive, negative, or neutral faces, or to pictures of places. For each type of trial, behavioral metrics include: total number of trials presented, number of correct responses, and accuracy (number of correct responses divided by the total number of trials). The mean and SD of the reaction times for correct responses are also included. Following the imaging session, the EN-Back Recognition Memory task asks the participants to decide if the pictures presented were previously seen in the EN-Back task. For each stimulus type (old and new), hit rates and false alarm rates are calculated. In addition, relevant metrics including corrected accuracy, response bias and d-prime are computed.

### Task-fMRI Analysis

Estimates of task-related activation strength are computed at the individual subject level using a general linear model (GLM) implemented in AFNI’s 3dDeconvolve (Cox, 1996). Preanalysis processing steps, which are not part of the fMRI preprocessing, include the removal of initial frames^3^ to ensure equilibration of the T_1_w signal and normalization of voxel time series by the mean across time of each voxel. Nuisance regressors are included to model the baseline and quadratic trends in the time-series data. Motion estimates and their derivatives are also included as regressors (Power et al., 2014). Time points with FD greater than 0.9 mm are censored (Siegel et al., 2014).

Prior to the use of motion estimates for regression and censoring, estimated motion time courses are temporally filtered using an infinite impulse response (IIR) notch filter, to attenuate signals in the range of 0.31 - 0.43 Hz. This frequency range corresponds to empirically observed oscillatory signals in the motion estimates that are linked to respiration and the dynamic changes in magnetic susceptibility due to movement of the lungs in the range of 18.6 - 25.7 respirations / minute. With the removal of these fast oscillations linked to respiration, the filtered motion estimates and FD values more accurately reflect actual head motion (Fair et al., 2018).

Hemodynamic response functions are modelled with two parameters using a gamma variate basis function plus its temporal derivative (using AFNI’s ‘SPMG’ option within 3dDeconvolve). The MID analysis model includes cue-elicited anticipation of large and small rewards or losses, or no incentive and feedback for large and small wins and losses. Linear contrasts are computed for anticipation of large and small reward vs. no incentive, anticipation of large and small loss vs. no incentive, feedback of win vs. missed win, and feedback of loss vs. avoided loss (Supp. Table 7). The SST model includes predictors for successful go trials (“correct go”), failed go trials (“incorrect go”), successful stop trials (“correct stop”), and failed stop trials (“incorrect stop”). Contrasts computed include correct go vs. fixation, correct stop vs. correct go, incorrect stop vs. correct go, any stop vs. correct go, correct stop vs. incorrect stop, incorrect go vs. correct go, and incorrect go vs. incorrect stop (Supp. Table 8). The EN-back model includes predictors for each type of stimulus (i.e., place and emotional face) in each of the EN-back conditions (i.e., 0-back and 2-back) plus fixation. Linear contrasts are obtained for 2-back vs. 0-back across stimulus types, emotional faces vs. places across memory loads, 2-back vs. 0-back for each stimulus type, and each memory load and each stimulus type vs. fixation (Supp. Table 9). For MID and SST analyses, events are modeled as instantaneous; for EN-back, the duration of cues (~3 s) and trial blocks (~24 s) are modeled as square waves convolved with the two parameter gamma basis function (i.e., block duration specified when using AFNI’s ‘SPMG’ option).

GLM beta coefficients and standard errors of the mean (SEM; calculated from the ratio of the beta and t-statistic) for voxels containing cortical gray matter are sampled onto the surface, projecting 1 mm from the gray/white boundary (“white” surface) into cortical gray matter along the surface normal vector at each cortical surface mesh point, or vertex (using FreeSurfer’s mri_vol2surf with “-projdist 1” option and default “nearest” interpolation). For each linear contrast specified for a given task, average coefficients and standard errors are calculated for cortical surface-based ROIs using FreeSurfer’s standard, anatomically-defined parcellation (Desikan et al., 2006), as well as a functionally-defined parcellation based on resting-state functional connectivity patterns (Gordon et al., 2014). Averages are also calculated for subcortical ROIs (Fischl et al., 2002). ROI average beta coefficients and standard errors are computed for each of two runs. We compute the average across runs for each participant weighted by the nominal degrees of freedom (number of frames remaining after motion censoring minus number of model parameters, but not accounting for temporal autocorrelation), which differs between runs due to motion censoring. Runs with fewer than 50 degrees of freedom are excluded from the average between runs.

The frequency and magnitude of head movements varies widely in children. Some participants exhibit frequent periods of motion resulting in a greatly reduced number of time points with which to estimate model parameters. Depending on when supra-threshold head movements (FD>0.9 mm) occur in relation to the instances of a given event type, rare conditions may be severely under-represented in some participants, or even lack representation entirely. For unrepresented conditions, beta and SEM values are undefined and shared as empty cells in the tabulated data. If conditions are under-represented, the design matrix of the GLM analysis becomes ill-conditioned, making the estimated beta weights unreliable for those event types and the contrasts that include them. In rare cases, this results in extreme values for the beta estimates, as much as several orders of magnitude different from typical beta values for a given contrast. The SEM is similarly increased in these cases. In the presence of extreme outliers, which violates standard parametric assumptions, group-level statistical analyses can produce invalid and nonsensical results. To prevent this, we censor the beta and SEM values if they are identified as having extremely high SEM values and therefore low reliability beta estimates. For a given subject with an extreme value for a particular contrast and ROI, there are typically outliers in other brain regions for the same subject and contrast and generally greater variation across brain regions. We censor the beta and SEM values for all ROIs for those contrasts that have root mean square (RMS) of SEM values across the brain greater than 5% signal change. This represents less than 0.5% of all subject-task-contrast-run combinations. The censored values are replaced with empty cells.

### Resting-State fMRI Analysis

Measures of functional connectivity are computed using a seed-based, correlational approach (Van Dijk et al., 2010), adapted for cortical surface-based analysis (Seibert and Brewer, 2011). Pre-analysis processing steps, which are not part of the fMRI preprocessing, include the removal of initial frames, normalization, regression, temporal filtering, and calculation of ROI-average time courses. After removing the initial frames^2^, we normalize voxel time series by the mean across time of each voxel and then use linear regression to remove quadratic trends, signals correlated with estimated motion time courses, and the mean time courses of cerebral white matter, ventricles, and whole brain, as well as their first derivatives (Power et al., 2014; Satterthwaite et al., 2012). The white matter, ventricle, and whole brain ROIs used to calculate mean time courses were derived from FreeSurfer’s automated brain segmentation (aseg), resampled into voxel-wise alignment with the fMRI data, and then eroded by a single fMRI-resolution voxel. Motion regression includes six parameters plus their derivatives and squares. Only frames with displacement (FD) below 0.3 mm are included in the regression (Power et al., 2014). This threshold is chosen only for the regression stage, and was viewed as a reasonable compromise, as ranges for FD thresholds for analysis stages might vary depending on the dataset. The analysis thresholds were set to FD < 0.2 mm (Power et al., 2014). After regression, time courses are band-pass filtered between 0.009 and 0.08 Hz (Hallquist et al., 2013). As described above for task fMRI analysis, estimated motion time courses are temporally filtered to attenuate signals linked to respiration.

Preprocessed time courses are sampled onto the cortical surface for each individual subject. Voxels containing cortical gray matter are sampled onto the surface in the same manner as the task-fMRI data. Average time courses are calculated for cortical surface-based ROIs using FreeSurfer’s standard, anatomically-defined parcellation (Desikan et al., 2006), as well as a functionally-defined parcellation based on resting-state functional connectivity patterns (Gordon et al., 2014), which are resampled from atlas-space to individual subject-space. Average time courses are also calculated for subcortical ROIs (Fischl et al., 2002). Variance across time is calculated for each ROI, a measure that reflects the magnitude of low frequency oscillations.

We calculate correlation values for each pair of ROIs, which are Fisher transformed to z-statistics and averaged within or between networks to provide summary measures of network correlation strength (Van Dijk et al., 2010). Within the Gordon parcellation, ROIs are grouped together into several networks (e.g., default, fronto-parietal, dorsal attention, etc.) (Gordon et al., 2014) (Supp. Table 4). Average correlation within a network is calculated as the average of the Fisher-transformed correlations for each unique, pairwise combination of ROIs belonging to that network. Average correlation between one network and another is calculated similarly by averaging the correlations for each unique, pair-wise combination of ROIs in the first network with the ROIs in the second. In addition, we calculate the correlation between each network and each subcortical gray matter ROI by averaging the correlations between each ROI in the network and a given subcortical ROI.

Motion censoring is used to reduce residual effects of head motion that may survive the pre-analysis regression (Power et al., 2012; Power et al., 2014). As noted above, time points with FD greater than 0.2 mm are excluded from the variance and correlation calculations. Time periods with fewer than five contiguous, sub-threshold time points are also excluded. The effects of head motion can potentially linger for several seconds after an abrupt head motion, for example due to spin-history or T1 relaxation effects (Friston et al., 1996), so an additional round of censoring is applied based on detecting time points that are outliers with respect to spatial variation across the brain. SD across ROIs is calculated for each time point, and outlier time points, defined as having an SD value more than three times the median absolute deviation (MAD) below or above the median SD value, are excluded from variance and correlation calculations.

### Quality Control

The DAIC in collaboration with ABCD partners are continually investigating ways to better curate these large datasets using enhanced QC procedures. This section is an overview of the QC procedures for ABCD Data Release 1.1. A detailed report of the ABCD QC procedures and an analysis of these QC metrics for the total baseline data (i.e., ABCD Data Release 2.0) is anticipated to be released in early 2019.

Using a combination of automated and manual methods, we review datasets for problems such as incorrect acquisition parameters, imaging artifacts, or corrupted data files. Automated protocol compliance checks are performed by the on-site FIONA workstations, providing feedback to the scan operators before upload to the DAIC about the completeness of the dataset and the adherence to the intended imaging parameters. After receipt of the data at the DAIC, protocol compliance information is recreated and uploaded to the ABCD REDCap Database. Out-of-compliance series are reviewed by DAIC staff, and sites are contacted if corrective action is required.

Protocol compliance criteria include whether key imaging parameters, such as voxel size or repetition time, match the expected values for a given scanner. For dMRI and fMRI series, the presence or absence of corresponding B0 distortion field map series is checked. Each imaging series is also checked for completeness to confirm that the number of files matches what was expected for each series on each scanner. Missing files are typically indicative of either an aborted scan or incomplete data transfer, of which the latter can usually be resolved through re-initiating the data transfer. Errors in the unpacking and processing of the imaging data at various stages are tracked, allowing for an assessment of the number of failures at each stage and prioritization of efforts to resolve problems and prevent future errors.

Automated quality control procedures include the calculation of metrics such as signal-to-noise ratio (SNR) and head motion statistics. For sMRI series, metrics include mean and SD of brain values. For dMRI series, head motion is estimated by registering each frame to a corresponding image synthesized from a tensor fit, accounting for variation in image contrast across diffusion orientations (Hagler et al., 2009). Overall head motion is quantified as the average of estimated FD. Dark slices, an artifact indicative of abrupt head motion, are identified as outliers in the RMS difference between the original data and data synthesized from tensor fitting. The total numbers of the slices and frames affected by these motion artifacts are calculated for each dMRI series. For fMRI series, measures include mean FD, or frame-to-frame head motion, the number of seconds with FD less than 0.2, 0.3, or 0.4 mm (Power et al., 2012), and temporal SNR (tSNR) (Triantafyllou et al., 2005) computed after motion correction.

Trained technicians visually review image series as part of our manual QC procedures, including T_1_w, T_2_w, dMRI, dMRI field maps, fMRI, and fMRI field maps. Reviewers inspect images for signs of artifacts and poor image quality, noting various imaging artifacts and flagging unacceptable data, typically those with the most severe artifacts or irregularities. Reviewers are shown several pre-rendered montages for each series, showing multiple slices and views of the first frame, and multiple frames of individual slices if applicable. For multi-frame images, linearly spaced subset of frames are shown as a 9 × 9 matrix of 81 frames. For dMRI and fMRI, derived images are also shown. For dMRI series, derived images include the average b=0 image, FA, MD, tensor fit residual error, and DEC FA map. For fMRI series, derived images include the average across time and the temporal SD (computed following motion correction).

All series are consensus rated by two or more reviewers. In the case of a rejection, the reviewer is required to provide notes indicating the types of artifacts observed using a standard set of abbreviations for commonly encountered artifacts. Series rejected based on data quality criteria are excluded from subsequent processing and analysis.

To ensure the quality of derived measures, trained technicians additionally review postprocessed sMRI data to evaluate the accuracy of cortical surface reconstruction. For each cortical surface reconstruction, reviewers gauge the severity of five categories of image artifact or reconstruction inaccuracy: motion, intensity inhomogeneity, white matter underestimation, pial overestimation, and magnetic susceptibility artifact. Numeric values are assigned on a scale of 0-3, indicating absent, mild, moderate, and severe levels of each type of artifact, respectively. The reviewers assign an overall QC score indicating whether the cortical surface reconstruction is recommended for use (1) or recommended for exclusion (0). Exclusion is recommended if any of the five categories are rated as severe (a value of 3).

For post-processed dMRI data, reviewers compare RSI-derived ND images (see *dMRI Microstructural Analysis)* to corresponding, co-registered T_1_w images, and rate each dMRI series along five dimensions of quality: residual B0 distortion, registration to the T_1_w image, image quality, segmentation integrity, and field of view (FOV) cutoff. For each, numeric values of 0-3 are assigned, indicating absent, mild, moderate, and severe. Residual distortion is assessed by looking for stretching or compression of white matter tracts in the ND image relative to the rigid-body co-registered T_1_w image, focusing on the corpus callosum and frontal lobe. Poor registration is rated on the basis of visible rotation or translation between the T_1_w and RSI-ND images. The image quality rating is based on the presence of banding, graininess, motion, artifacts, or poor gray/white contrast in the ND image. The automatic white matter tract segmentation is assessed for incompleteness, absence, or gross mis-location. FOV cutoff indicates clipping of the dorsal or ventral aspect of the cortex. Each dMRI series is then assigned an overall QC score of recommended for use (1) or recommended for exclusion (0). A series will be recommended for exclusion (QC score of 0) if B_0_ warp, registration, image quality, or segmentation are rated as severe (a value of 3). While FOV cutoff is assessed, it is not used as a factor in deciding the overall QC score.

For ABCD Data Release 1.1, only automated and manual QC of fMRI data was performed as detailed above. The DAIC and partners are currently investigating optimal postprocessed fMRI reviews, which would resemble in part the dMRI QC procedure. Further details will be provided with the full baseline release in ABCD Data Release 2.0.

### Data Sharing

Public sharing of unprocessed, preprocessed, and derived imaging data is undertaken in partnership with the NDA. We have implemented methods and procedures for sharing of imaging data in the following three forms: DICOM files, preprocessed NIfTI files, and tabulated results of ROI-based analyses for each modality. Starting April 2017, DICOM files were made publicly available via a Fast Track mechanism. These DICOM files are released on NDA (https://data-archive.nimh.nih.gov/abcd) on a continual basis within approximately one month of data collection using the ABCD fast-track image sharing scripts (https://scicrunch.org/resolver/SCR_016021). DICOM files are arranged in BIDS directory format with folders of DICOM images per image series packaged in individual archive files (tgz) for each series. Metadata are included in the form of JSON-format text files, and task-fMRI series also include files containing stimulus and behavioral response timing information exported from the stimulus program (E-prime). A copy of the meta data is uploaded to NDA’s ‘image03’ datatype to link information to non-imaging-based assessments for the same participants.

In addition to the Fast Track data sharing, there are curated data releases that include results derived from imaging data using the processing pipeline described herein. The first of these curated data releases (ABCD Data Release 1.0) was made public on February 12, 2018 (https://data-archive.nimh.nih.gov/abcd). It is considered an “interim” release because it includes slightly less than half of the projected baseline sample. The second annual release (ABCD Data Release 2.0) includes the complete 2016-2018 baseline sample and is scheduled for mid-2019. As participants return every two years for successive waves of imaging, subsequent releases will include longitudinal data. Mid-year “patch” releases (e.g., ABCD Data Release 1.1) may also be shared as appropriate to address minor issues or salvage previously missing data. Descriptions of the specific changes between successive data releases will be included in comprehensive data release notes provided with each release. This manuscript documents the methods and data shared as part of the ABCD Data Release 1.1, available November 2018.

Preprocessed imaging data, as described in the “preprocessing” sections above, are packaged in archive files (tgz) for each image series containing BIDS formatted directory trees and NIfTI format data files (software to share preprocessed data: https://scicrunch.org/resolver/SCR_016016; consistent with BIDS specifications version 1.1.1: http://bids.neuroimaging.io/bids_spec.pdf). Imaging metadata derived from the original DICOM files are packaged along with each preprocessed data series as JSON files. Additional dMRI-specific information included diffusion gradients adjusted for head rotation (bvecs.txt), diffusion gradient strengths (bvals.txt), and a rigid-body transformation matrix specifying the registration between the dMRI image and the corresponding processed sMRI T_1_w image (stored in the JSON file). Additional fMRI-specific information includes estimated motion time courses and a rigid-body transformation matrix specifying the registration between the fMRI image and the T_1_w image (stored in the JSON file). For task-fMRI series, event timing information is included as tab-separated value (tsv) files.

The results of additional processing and ROI analysis are shared in tabulated form to the NDA database (https://scicrunch.org/resolver/SCR_016010), from which users can export spreadsheet files (tsv). Tabulated results derived from sMRI image analysis include morphometrics and image intensity measures, as well as quality control measures for FreeSurfer cortical surface reconstruction. Measures derived from dMRI include the results of DTI and RSI analyses, providing information about brain tissue microstructure. Resting-state fMRI-derived tabulated results include correlations within and between pre-defined cortical networks, average correlation between each network and each subcortical ROI, and the low frequency BOLD signal variance in each subcortical ROI (Fischl et al., 2002), Gordon parcel (Gordon et al., 2014), and standard FreeSurfer parcellation (Desikan et al., 2006). Also included are mean motion measures and the number of fMRI time points (TRs) before and after censoring of frames with excessive motion. For task-fMRI, beta and SEM values are averaged within ROIs and tabulated for each contrast for a given task, for run 1, run 2, and the average across runs. Also included are mean motion measures, the number of TRs before and after censoring, the number of degrees of freedom, and a comprehensive set of behavioral performance measures.

### Methods Sharing

Because faithfully reproducing complex image processing methods and workflows based on published descriptions alone has become increasingly difficult, public sharing of methods is an integral part of the data sharing mission of the ABCD Study. We have packaged the processing pipeline described here within a portable container format (Docker container), which includes a lightweight, virtualized Linux environment, and the complete execution environment, including all required software dependencies. This stand-alone, platform-independent executable can be used in any Docker-enabled processing environment and is specifically designed to work with ABCD DICOM data shared via the Fast Track data sharing mechanism. Packaging the processing pipeline in this way eliminates the need to install third-party processing tools and libraries, enabling easy and generic installation on all systems and removing hidden dependencies, and guarantees standardized processing workflows. For tools written with MATLAB, including those contained within MMPS, MATLAB functions were compiled and packaged with the MATLAB Compiler Runtime library. Because the executable Docker container contains third party software requiring licenses (i.e., FreeSurfer), users must supply a license file to be embedded in the local copy of the Docker container. Users may currently download a beta-testing version of the executable Docker container from NITRC (https://www.nitrc.org/projects/abcd_study). In the future, these tools will be made available through NDA with execution in the Amazon cloud. Also, a “Dockerfile”, which is a text file specifying the contents of a Docker container, will be made available via GitHub (github.com/ABCD-STUDY), enabling automated download and installation of the software components.

## Discussion

The ABCD Study will provide the most comprehensive longitudinal investigation to date of the neurobiological trajectories of brain and behavior development from late childhood through adolescence to early adulthood. There are many risk processes during adolescence that lead to chronic diseases in later life, including tobacco use, alcohol and illicit substance use, unsafe sex, obesity, sports injury and lack of physical activity (Patton et al., 2017). Furthermore, this developmental period can give rise to many common psychiatric conditions including anxiety disorders, bipolar disorder, depression, eating disorders, psychosis, and substance abuse (Hafner et al., 1989; Kessler et al., 2005). The ABCD Study is well-positioned to capture the behavioral and neurobiological changes taking place during healthy development, prodromal behavioral issues and antecedent neuroanatomical changes. Identifying neuropsychological, structural and functional measures as potential biomarkers of disorders may better inform the diagnosis and treatment of youth who present with early mental and physical health concerns.

Rigorous acquisition monitoring and uniform image processing of a large cohort of ethnically diverse young people regularly over adolescence is necessary to characterize subtle neuroanatomical and functional changes during such a plastic developmental period. Curated data releases providing canonical neuroimaging measures for this large dataset will enable the scientific community to test innumerable hypotheses related to brain and cognitive development. Meanwhile, the DAIC and ABCD collaborators will continue working to improve and extend the image processing pipeline, resulting in new and evolving imaging metrics to be evaluated for potential inclusion in future, curated data releases. Considering these novel metrics alongside canonical measurements provides the ability to compare and contrast new and existing analytical techniques. The capability to regularly reprocess all data for new analytic pipelines is made possible by the framework described in this manuscript. The intent is for the methods developed here to be shareable and deployable for other large scale neuroimaging studies.

### Limitations and Caveats

On an ongoing basis, the Fast Track data sharing mechanism occurs shortly after data collection, without processing, quality control, or curation, and includes all ABCD imaging data available and permitted to be shared. Imaging series with the most severe artifacts or those with missing or corrupted DICOM files are excluded from subsequent processing, and so are not included in the preprocessed or tabulated data sharing. For a given modality, additional participants may be missing from the tabulated data due to failures in brain segmentation or modality-specific processing and analysis. However, successfully processed data with moderate imaging artifacts are included in the preprocessed and tabulated data sharing. This is necessary to enable certain studies, such as methodological investigations of the effect of imaging artifacts on derived measures. For most group analyses, we recommend excluding cases with significant incidental findings, excessive motion, or other artifacts, and we provide a variety of QC-related metrics for use in applying sets of modality-specific, inclusion criteria (Supplemental Table 10; see *NDA 1.1 Release Notes ABCD Imaging Instruments* for additional details). Some researchers may wish to use this as a template for customized inclusion criteria that could include additional QC metrics or apply more conservative thresholds.

While Fast Track provides early access to DICOM data, this comes with the caveat that users may process the data inappropriately, resulting in inaccurate or spurious findings. The purpose of the ABCD processing pipeline is to provide images and derived measures for curated releases using consensus-derived approaches for processing and analysis. We suggest that authors clearly state the version of the curated release used (e.g., DOI: 10.15154/1460410, ABCD Data Release 1.1, November 2018), as the processing pipeline is expected to change over time. Each new release will document changes and will include data provided in previous releases, reprocessed as necessary to maintain consistency within a particular release. Combining data between curated releases is not recommended.

### Future Directions

Mirroring the general progression of neuroimaging processing tools and analysis methods in the field, the processing pipeline used for future ABCD data releases is expected to evolve over time. Other image analysis methods, either recently developed or soon to be developed, may better address particular issues, perhaps by providing superior correction of imaging artifacts, more accurate brain segmentation, or additional types of biologically relevant, imaging-derived measures. Alternative approaches for a given stage of processing or analysis will be compared using quantitative metrics of the reliability of the derived results. We seek to enhance and augment ABCD image analysis over time by incorporating new methods and approaches as they are implemented and validated.

To enable researchers to more easily take advantage of this data resource, the DAIC is providing a web-based tool called the ABCD Data Exploration and Analysis Portal (DEAP, part of the ABCD Data Release 2.0). This tool provides convenient access to the complete battery of ROI-based, multimodal imaging-derived measures as well as sophisticated statistical routines using generalized additive mixed models to analyze repeated measures and appropriately model the effects of site, scanner, family relatedness, and a range of demographic variables. Future releases will also support voxel-wise (volumetric) and vertex-wise (surface-based) analyses.

As a ten-year longitudinal study with participants returning every two years for successive waves of imaging, future data releases will include multiple time points for each participant. All of the imaging-derived measures provided for the baseline time point will also be generated independently for each successive time point. In addition, within-subject, longitudinal analyses will provide more sensitive estimates of longitudinal change for some imaging measures. For example, the FreeSurfer longitudinal processing stream reduces variability and increases sensitivity in the measurement of changes in cortical thickness and the volumes of subcortical structures. It does this by creating unbiased, within-subject templates and reinitializing surface reconstruction and brain segmentation using consensus information (Reuter and Fischl, 2011; Reuter et al., 2010; Reuter et al., 2012). With this approach, the within-subject template would be recreated with each new time point and all longitudinal change estimates recomputed. Analyses of cortical and subcortical changes in volume with even greater sensitivity is available through Quantitative Analysis of Regional Change (QUARC) (Holland et al., 2009; Holland and Dale, 2011; Holland et al., 2012; Thompson and Holland, 2011). A diffusion-based variant of QUARC is also available for longitudinal analysis of dMRI data with improved sensitivity (Holland and Dale, 2011; McDonald et al., 2010). The goal of future development and testing will be to provide a collection of longitudinal change estimates for a variety of imaging-derived measures that are maximally sensitive to small changes, are unbiased with respect to the ordering of time points, and have a slope of one with respect to change estimated from time points processed independently.

An important factor when analyzing multi-site and longitudinal imaging data is ensuring the comparability of images between scanners, referred to as data harmonization. Statistical harmonization procedures are presently available to correct for variations between scanners. For example, the unique identifier for each scanner (i.e., DeviceSerialNumber) can be used as a categorical covariate to account for potential differences between individual scanners in the mean value of a given measure (Brown et al., 2012). Recently, a genomic batch-effect correction tool has been proposed as a tool for data harmonization across multiple scanners for measures of cortical thickness (Fortin et al., 2017a) and diffusivity (Fortin et al., 2017b). This procedure can be applied to curated data releases by individual researchers or incorporated into the statistical manipulation front-end tool (DEAP, https://scicrunch.org/resolver/SCR_016158) if considered reliable and robust. Another harmonization approach attempts to retain site-specific features by generating study-specific atlases per scanner, which was shown to yield modest improvements in a longitudinal aging investigation (Erus et al., 2018). However, that particular study-specific approach is technically complicated and processor intensive for replication. For diffusion imaging, one promising approach uses rotation invariant spherical harmonics within a multimodal image registration framework (Mirzaalian et al., 2018). The DAIC is actively testing new approaches for harmonization of images prior to segmentation, parcellation, and DTI/RSI fitting to improve compatibility across sites and time. Over the coming years, the ABCD Study will incorporate refinements to these harmonization techniques and reprocess data from prior releases to reduce between-scanner differences.

### Conclusion

We have described the processing pipeline being used to generate the preprocessed imaging data and derived measures that were included in the ABCD Data Release made available October 2018 in a public data release through NDA (DOI: 10.15154/1460410, ABCD Data Release 1.1, October 2018; see https://data-archive.nimh.nih.gov/abcd). This resource includes multimodal imaging data, a comprehensive battery of behavioral assessments, and demographic information, on a group of 4,521 typically developing children between the ages of 9 and 10. Neuroimaging-derived measures include morphometry, microstructure, functional associations, and task-related functional activations. This is the first of two years of baseline recruitment for this ten-year, longitudinal study of brain development and peri-adolescent health. Data collection for the second year of baseline recruitment will be completed in late 2018. Processed data and derived measures for the entire baseline sample, projected to be approximately 11,500 participants, will be included in ABCD Data Release 2.0, which will be made available mid-2019. When complete, the ABCD dataset will provide a remarkable opportunity to comprehensively study the relationships between brain and cognitive development, substance use and other experiences, and social, genetic, and environmental factors. It will allow the scientific community to address many important questions about brain and behavioral development and about the genetic architecture of neural and other behaviorally relevant phenotypes. The processing pipeline we have described provides a comprehensive battery of multimodal imaging-derived measures. The processing methods include corrections for various distortions, head motion in dMRI and fMRI images, and intensity inhomogeneity in structural images. Collectively, these corrections are designed to reduce variance of estimated change (Holland et al., 2010), increase the accuracy of registration between modalities, and improve the accuracy of brain segmentation and cortical surface reconstruction.

## Footnotes

1 Whether a participant performed the MID, SST, or EN-back task in the scanner or on a laptop is indicated respectively by the following variables: ra_scan_cl_mid_scan_lap, ra_scan_cl_nbac_scan_lap, ra_scan_cl_sst_scan_lap. For more details, see “ABCD RA Scanning Checklist and Notes” available at https://ndar.nih.gov (instrument ‘abcd_ra’).

2 Offline reconstruction of multiband data is required for GE scanners with software version DV25. Starting in September 2017, GE scanners at three ABCD sites were upgraded to software version DV26, supporting online reconstruction of multiband data and providing DICOM files. The remaining GE scanners at ABCD sites were upgraded to DV26 in September 2018.

3 A total of 16 initial frames (12.8 seconds) are discarded. On Siemens and Philips scanners, the first eight frames make up the pre-scan reference, and are not saved as DICOMS. An additional eight frames are discarded as part of the pre-analysis processing, for a total of 16 initial frames. On GE scanners with software version DV25, the first 12 frames make up the pre-scan reference. Instead of being discarded, those 12 reference scans are combined into one, and saved as the first frame, for a total of five initial frames to be discarded as part of the pre-analysis processing for GE DV25 series. On GE scanners with software version DV26, the pre-scan reference is not retained at all, and a total of 16 initial frames are discarded for GE DV26 scans as part of the pre-analysis processing.

## Acknowledgements

Data used in the preparation of this article were obtained from the Adolescent Brain Cognitive Development (ABCD) Study (https://abcdstudy.org), held in the NIMH Data Archive (NDA). This is a multisite, longitudinal study designed to recruit more than 10,000 children aged 9-10 years and follow them over 10 years into early adulthood. The ABCD Study is supported by the National Institutes of Health and additional federal partners under award numbers U01DA041022, U01DA041028, U01DA041048, U01DA041089, U01DA041106, U01DA041117, U01DA041120, U01DA041134, U01DA041148, U01DA041156, U01DA041174, U24DA041123, and U24DA041147. A full list of supporters is available at https://abcdstudy.org/nih-collaborators. A listing of participating sites and a complete listing of the study investigators can be found at https://abcdstudy.org/principal-investigators.html. ABCD consortium investigators designed and implemented the study and/or provided data but did not necessarily participate in analysis or writing of this report. This manuscript reflects the views of the authors and may not reflect the opinions or views of the NIH or ABCD consortium investigators. The ABCD data repository grows and changes over time. The ABCD data used in this report came from NIMH Data DOI: 10.15154/1460410.

Representatives from NIH (NIDA, NIMH, NIAAA, NIMHD,OBSSR, ORWH) contributed to the interpretation of the data and participated in the preparation, review and approval of the manuscript. The views and opinions expressed in this manuscript are those of the authors only and do not necessarily represent the views, official policy or position of the U.S. Department of Health and Human Services or any of its affiliated institutions or agencies.

## Disclosure Statement

Martin Paulus is an adviser to Spring Care (New York City, NY, USA), a behavioral health startup. He has received royalties for an article about methamphetamine in UpToDate (Wolters Kluwer; Waltham, MA, USA), and received grant support from Janssen Pharmaceuticals.

W. Kyle Simmons is an employee of Janssen Research and Development, LLC.

Andrew Nencka receives research funding from GE Healthcare.

Kevin M. Gray provided consultation for Pfizer, Inc.

Susan R.B. Weiss and her husband own stock in Merck and GE, respectively.

Anders M. Dale reports that he is a Founder of and holds equity in CorTechs Labs, Inc., and serves on its Scientific Advisory Board. He is a member of the Scientific Advisory Board of Human Longevity, Inc., and receives funding through research grants with General Electric Healthcare. The terms of these arrangements have been reviewed by and approved by the University of California, San Diego in accordance with its conflict of interest policies.

All other authors report no potential conflicts of interest.

## References Cited

Alexander, A.L., Lee, J.E., Lazar, M., Field, A.S., 2007. Diffusion tensor imaging of the brain. Neurotherapeutics 4, 316–329.

Andersson, J.L., Skare, S., Ashburner, J., 2003. How to correct susceptibility distortions in spinecho echo-planar images: application to diffusion tensor imaging. Neuroimage 20, 870–888.

Andersson, J.L.R., Sotiropoulos, S.N., 2016. An integrated approach to correction for off-resonance effects and subject movement in diffusion MR imaging. Neuroimage 125, 1063–1078.

Ashburner, J., Friston, K.J., 2000. Voxel-based morphometry--the methods. Neuroimage 11, 805–821.

Bagot, K.S., Matthews, S.A., Mason, M., Squeglia, L.M., Fowler, J., Gray, K., Herting, M., May, A., Colrain, I., Godino, J., Tapert, S., Brown, S., Patrick, K., 2018. Current, future and potential use of mobile and wearable technologies and social media data in the ABCD study to increase understanding of contributors to child health. Dev Cogn Neurosci 32, 121–129.

Barch, D.M., Albaugh, M.D., Avenevoli, S., Chang, L., Clark, D.B., Glantz, M.D., Hudziak, J.J., Jernigan, T.L., Tapert, S.F., Yurgelun-Todd, D., Alia-Klein, N., Potter, A.S., Paulus, M.P., Prouty, D., Zucker, R.A., Sher, K.J., 2018. Demographic, physical and mental health assessments in the adolescent brain and cognitive development study: Rationale and description. Dev Cogn Neurosci 32, 55–66.

Barnett, A.S., Hutchinson, E., Irfanoglu, M.O., Pierpaoli, C., 2014. Higher order correction of eddy current distortion in diffusion weighted echo planar images., Join Annual Meeting ISMRM-ESMRMB, Milan, Italy, p. 5119.

Basser, P.J., Mattiello, J., LeBihan, D., 1994a. Estimation of the effective self-diffusion tensor from the NMR spin echo. J Magn Reson B 103, 247–254.

Basser, P.J., Mattiello, J., LeBihan, D., 1994b. MR diffusion tensor spectroscopy and imaging. Biophys J 66, 259–267.

Basser, P.J., Pierpaoli, C., 1996. Microstructural and physiological features of tissues elucidated by quantitative-diffusion-tensor MRI. J Magn Reson B 111, 209–219.

Brown, T.T., Kuperman, J.M., Chung, Y., Erhart, M., McCabe, C., Hagler, D.J., Jr., Venkatraman, V.K., Akshoomoff, N., Amaral, D.G., Bloss, C.S., Casey, B.J., Chang, L., Ernst, T.M., Frazier, J.A., Gruen, J.R., Kaufmann, W.E., Kenet, T., Kennedy, D.N., Murray, S.S., Sowell, E.R., Jernigan, T.L., Dale, A.M., 2012. Neuroanatomical assessment of biological maturity. Curr Biol 22, 1693–1698.

Brown, T.T., Kuperman, J.M., Erhart, M., White, N.S., Roddey, J.C., Shankaranarayanan, A., Han, E.T., Rettmann, D., Dale, A.M., 2010. Prospective motion correction of high-resolution magnetic resonance imaging data in children. Neuroimage 53, 139–145.

Casey, B.J., Cannonier, T., Conley, M.I., Cohen, A.O., Barch, D.M., Heitzeg, M.M., Soules, M.E., Teslovich, T., Dellarco, D.V., Garavan, H., Orr, C.A., Wager, T.D., Banich, M.T., Speer, N.K., Sutherland, M.T., Riedel, M.C., Dick, A.S., Bjork, J.M., Thomas, K.M., Chaarani, B., Mejia, M.H., Hagler, D.J., Jr., Daniela Cornejo, M., Sicat, C.S., Harms, M.P., Dosenbach, N.U.F., Rosenberg, M., Earl, E., Bartsch, H., Watts, R., Polimeni, J.R., Kuperman, J.M., Fair, D.A., Dale, A.M., Workgroup, A.I.A., 2018. The Adolescent Brain Cognitive Development (ABCD) study: Imaging acquisition across 21 sites. Dev Cogn Neurosci 32, 43–54.

Chang, H., Fitzpatrick, J.M., 1992. A technique for accurate magnetic resonance imaging in the presenceof field inhomogeneities. IEEE Trans Med Imaging 11, 319–329.

Chen, C.H., Gutierrez, E.D., Thompson, W., Panizzon, M.S., Jernigan, T.L., Eyler, L.T., Fennema-Notestine, C., Jak, A.J., Neale, M.C., Franz, C.E., Lyons, M.J., Grant, M.D., Fischl, B., Seidman, L.J., Tsuang, M.T., Kremen, W.S., Dale, A.M., 2012. Hierarchical genetic organization of human cortical surface area. Science 335, 1634–1636.

Cohen, A.O., Conley, M.I., Dellarco, D.V., Casey, B.J., 2016. The impact of emotional cues on short-term and long-term memory during adolescence. Program No. 90.25 Neuroscience Meeting Planner., San Diego, CA: Society for Neuroscience 2016. Online.

Cox, R.W., 1996. AFNI: software for analysis and visualization of functional magnetic resonance neuroimages. Comput Biomed Res 29, 162–173.

Dale, A.M., Fischl, B., Sereno, M.I., 1999. Cortical surface-based analysis. I. Segmentation and surface reconstruction. Neuroimage 9, 179–194.

Dale, A.M., Sereno, M.I., 1993. Improved Localizadon of Cortical Activity by Combining EEG and MEG with MRI Cortical Surface Reconstruction: A Linear Approach. J Cogn Neurosci 5, 162–176.

Desikan, R.S., Segonne, F., Fischl, B., Quinn, B.T., Dickerson, B.C., Blacker, D., Buckner, R.L., Dale, A.M., Maguire, R.P., Hyman, B.T., Albert, M.S., Killiany, R.J., 2006. An automated labeling system for subdividing the human cerebral cortex on MRI scans into gyral based regions of interest. Neuroimage 31, 968–980.

Dosenbach, N.U.F., Koller, J.M., Earl, E.A., Miranda-Dominguez, O., Klein, R.L., Van, A.N., Snyder, A.Z., Nagel, B.J., Nigg, J.T., Nguyen, A.L., Wesevich, V., Greene, D.J., Fair, D.A., 2017. Real-time motion analytics during brain MRI improve data quality and reduce costs. Neuroimage 161, 80–93.

Elman, J.A., Panizzon, M.S., Hagler, D.J., Jr., Fennema-Notestine, C., Eyler, L.T., Gillespie, N.A., Neale, M.C., Lyons, M.J., Franz, C.E., McEvoy, L.K., Dale, A.M., Kremen, W.S., 2017. Genetic and environmental influences on cortical mean diffusivity. Neuroimage 146, 90–99.

Erus, G., Doshi, J., An, Y., Verganelakis, D., Resnick, S.M., Davatzikos, C., 2018. Longitudinally and inter-site consistent multi-atlas based parcellation of brain anatomy using harmonized atlases. Neuroimage 166, 71–78.

Fair, D.A., Miranda-Dominguez, O., Snyder, A.Z., Perrone, A.A., Earl, E.A., Van, A.N., Koller, J.M., Feczko, E., Klein, R.L., Mirro, A.E., Hampton, J.M., Adeyemo, B., Laumann, T.O., Gratton, C., Greene, D.J., Schlaggar, B., Hagler, D., Watts, R., Garavan, H., Barch, D.M., Nigg, J.T., Petersen, S.E., Dale, A., Feldstein-Ewing, S.W., Nagel, B.J., Dosenbach, N.U.F., 2018. Correction of respiratory artifacts in MRI head motion estimates. bioRxiv.

Fair, D.A., Nigg, J.T., Iyer, S., Bathula, D., Mills, K.L., Dosenbach, N.U., Schlaggar, B.L., Mennes, M., Gutman, D., Bangaru, S., Buitelaar, J.K., Dickstein, D.P., Di Martino, A., Kennedy, D. N., Kelly, C., Luna, B., Schweitzer, J.B., Velanova, K., Wang, Y.F., Mostofsky, S., Castellanos, F.X., Milham, M.P., 2012. Distinct neural signatures detected for ADHD subtypes after controlling for micro-movements in resting state functional connectivity MRI data. Front Syst Neurosci 6, 80.

Fischl, B., 2012. FreeSurfer. Neuroimage 62, 774–781.

Fischl, B., Dale, A.M., 2000. Measuring the thickness of the human cerebral cortex from magnetic resonance images. Proc Natl Acad Sci U S A 97, 11050–11055.

Fischl, B., Liu, A., Dale, A.M., 2001. Automated manifold surgery: constructing geometrically accurate and topologically correct models of the human cerebral cortex. IEEE Trans Med Imaging 20, 70–80.

Fischl, B., Salat, D.H., Busa, E., Albert, M., Dieterich, M., Haselgrove, C., van der Kouwe, A., Killiany, R., Kennedy, D., Klaveness, S., Montillo, A., Makris, N., Rosen, B., Dale, A.M., 2002. Whole brain segmentation: automated labeling of neuroanatomical structures in the human brain. Neuron 33, 341–355.

Fischl, B., Sereno, M.I., Dale, A.M., 1999a. Cortical surface-based analysis. II: Inflation, flattening, and a surface-based coordinate system. Neuroimage 9, 195–207.

Fischl, B., Sereno, M.I., Tootell, R.B., Dale, A.M., 1999b. High-resolution intersubject averaging and a coordinate system for the cortical surface. Hum Brain Mapp 8, 272–284.

Fischl, B., van der Kouwe, A., Destrieux, C., Halgren, E., Segonne, F., Salat, D.H., Busa, E., Seidman, L.J., Goldstein, J., Kennedy, D., Caviness, V., Makris, N., Rosen, B., Dale, A.M., 2004. Automatically parcellating the human cerebral cortex. Cereb Cortex 14, 11–22.

Fortin, J.P., Cullen, N., Sheline, Y.I., Taylor, W.D., Aselcioglu, I., Cook, P.A., Adams, P., Cooper, C., Fava, M., McGrath, P.J., McInnis, M., Phillips, M.L., Trivedi, M.H., Weissman, M.M., Shinohara, R.T., 2017a. Harmonization of cortical thickness measurements across scanners and sites. Neuroimage 167, 104–120.

Fortin, J.P., Parker, D., Tunc, B., Watanabe, T., Elliott, M.A., Ruparel, K., Roalf, D.R., Satterthwaite, T.D., Gur, R.C., Gur, R.E., Schultz, R.T., Verma, R., Shinohara, R.T., 2017b. Harmonization of multi-site diffusion tensor imaging data. Neuroimage 161, 149–170.

Friston, K.J., Ashburner, J., Frith, C.D., J-B., P., J.D., H., Frackowiak, R.S.J., 1995. Spatial registration and normalization of images. Hum Brain Mapp 3, 165–189.

Friston, K.J., Williams, S., Howard, R., Frackowiak, R.S., Turner, R., 1996. Movement-related effects in fMRI time-series. Magn Reson Med 35, 346–355.

Garavan, H., Bartsch, H., Conway, K., Decastro, A., Goldstein, R.Z., Heeringa, S., Jernigan, T., Potter, A., Thompson, W., Zahs, D., 2018. Recruiting the ABCD sample: Design considerations and procedures. Dev Cogn Neurosci 32, 16–22.

Ghosh, S.S., Kakunoori, S., Augustinack, J., Nieto-Castanon, A., Kovelman, I., Gaab, N., Christodoulou, J.A., Triantafyllou, C., Gabrieli, J.D., Fischl, B., 2010. Evaluating the validity of volume-based and surface-based brain image registration for developmental cognitive neuroscience studies in children 4 to 11 years of age. Neuroimage 53, 85–93.

Glasser, M.F., Smith, S.M., Marcus, D.S., Andersson, J.L., Auerbach, E.J., Behrens, T.E., Coalson, T.S., Harms, M.P., Jenkinson, M., Moeller, S., Robinson, E.C., Sotiropoulos, S.N., Xu, J., Yacoub, E., Ugurbil, K., Van Essen, D.C., 2016. The Human Connectome Project’s neuroimaging approach. Nat Neurosci 19, 1175–1187.

Gordon, E.M., Laumann, T.O., Adeyemo, B., Huckins, J.F., Kelley, W.M., Petersen, S.E., 2014. Generation and Evaluation of a Cortical Area Parcellation from Resting-State Correlations. Cereb Cortex.

Govindan, R.M., Asano, E., Juhasz, C., Jeong, J.W., Chugani, H.T., 2013. Surface-based laminar analysis of diffusion abnormalities in cortical and white matter layers in neocortical epilepsy. Epilepsia 54, 667–677.

Hafner, H., Riecher, A., Maurer, K., Loffler, W., Munk-Jorgensen, P., Stromgren, E., 1989. How does gender influence age at first hospitalization for schizophrenia? A transnational case register study. Psychol Med 19, 903–918.

Hagler, D.J., Jr., Ahmadi, M.E., Kuperman, J., Holland, D., McDonald, C.R., Halgren, E., Dale, A.M., 2009. Automated white-matter tractography using a probabilistic diffusion tensor atlas: Application to temporal lobe epilepsy. Hum Brain Mapp 30, 1535–1547.

Hallquist, M.N., Hwang, K., Luna, B., 2013. The nuisance of nuisance regression: spectral misspecification in a common approach to resting-state fMRI preprocessing reintroduces noise and obscures functional connectivity. Neuroimage 82, 208–225.

Holland, D., Brewer, J.B., Hagler, D.J., Fennema-Notestine, C., Dale, A.M., 2009. Subregional neuroanatomical change as a biomarker for Alzheimer’s disease. Proc Natl Acad Sci U S A 106, 20954–20959.

Holland, D., Dale, A.M., 2011. Nonlinear registration of longitudinal images and measurement of change in regions of interest. Med Image Anal 15, 489–497.

Holland, D., Kuperman, J.M., Dale, A.M., 2010. Efficient correction of inhomogeneous static magnetic field-induced distortion in Echo Planar Imaging. Neuroimage 50, 175–183.

Holland, D., McEvoy, L.K., Dale, A.M., 2012. Unbiased comparison of sample size estimates from longitudinal structural measures in ADNI. Hum Brain Mapp 33, 2586–2602.

Jenkinson, M., Beckmann, C.F., Behrens, T.E., Woolrich, M.W., Smith, S.M., 2012. Fsl. Neuroimage 62, 782–790.

Jernigan, T.L., Brown, S.A., Coordinators, A.C., 2018. Introduction. Dev Cogn Neurosci 32, 1–3.

Jernigan, T.L., Brown, T.T., Hagler, D.J., Jr., Akshoomoff, N., Bartsch, H., Newman, E., Thompson, W.K., Bloss, C.S., Murray, S.S., Schork, N., Kennedy, D.N., Kuperman, J.M., McCabe, C., Chung, Y., Libiger, O., Maddox, M., Casey, B.J., Chang, L., Ernst, T.M., Frazier, J.A., Gruen, J.R., Sowell, E.R., Kenet, T., Kaufmann, W.E., Mostofsky, S., Amaral, D.G., Dale, A.M., Pediatric Imaging, N., Genetics, S., 2016. The Pediatric Imaging, Neurocognition, and Genetics (PING) Data Repository. Neuroimage 124, 1149–1154.

Jovicich, J., Czanner, S., Greve, D., Haley, E., van der Kouwe, A., Gollub, R., Kennedy, D., Schmitt, F., Brown, G., Macfall, J., Fischl, B., Dale, A., 2006. Reliability in multi-site structural MRI studies: effects of gradient non-linearity correction on phantom and human data. Neuroimage 30, 436–443.

Joyner, A.H., J, C.R., Bloss, C.S., Bakken, T.E., Rimol, L.M., Melle, I., Agartz, I., Djurovic, S., Topol, E.J., Schork, N.J., Andreassen, O.A., Dale, A.M., 2009. A common MECP2 haplotype associates with reduced cortical surface area in humans in two independent populations. Proc Natl Acad Sci U S A 106, 15483–15488.

Kang, X., Herron, T.J., Turken, A.U., Woods, D.L., 2012. Diffusion properties of cortical and pericortical tissue: regional variations, reliability and methodological issues. Magn Reson Imaging 30, 1111–1122.

Kessler, R.C., Berglund, P., Demler, O., Jin, R., Merikangas, K.R., Walters, E.E., 2005. Lifetime prevalence and age-of-onset distributions of DSM-IV disorders in the National Comorbidity Survey Replication. Arch Gen Psychiatry 62, 593–602.

Knutson, B., Westdorp, A., Kaiser, E., Hommer, D., 2000. FMRI visualization of brain activity during a monetary incentive delay task. Neuroimage 12, 20–27.

Kremen, W.S., Prom-Wormley, E., Panizzon, M.S., Eyler, L.T., Fischl, B., Neale, M.C., Franz, C. E., Lyons, M.J., Pacheco, J., Perry, M.E., Stevens, A., Schmitt, J.E., Grant, M.D., Seidman, L.J., Thermenos, H.W., Tsuang, M.T., Eisen, S.A., Dale, A.M., Fennema-Notestine, C., 2010. Genetic and environmental influences on the size of specific brain regions in midlife: the VETSA MRI study. Neuroimage 49, 1213–1223.

Kuperman, J.M., Brown, T.T., Ahmadi, M.E., Erhart, M.J., White, N.S., Roddey, J.C., Shankaranarayanan, A., Han, E.T., Rettmann, D., Dale, A.M., 2011. Prospective motion correction improves diagnostic utility of pediatric MRI scans. Pediatr Radiol 41, 1578–1582.

Leemans, A., Jones, D.K., 2009. The B-matrix must be rotated when correcting for subject motion in DTI data. Magn Reson Med 61, 1336–1349.

Levman, J., MacDonald, P., Lim, A.R., Forgeron, C., Takahashi, E., 2017. A pediatric structural MRI analysis of healthy brain development from newborns to young adults. Hum Brain Mapp 38, 5931–5942.

Lisdahl, K.M., Sher, K.J., Conway, K.P., Gonzalez, R., Feldstein Ewing, S.W., Nixon, S.J., Tapert, S., Bartsch, H., Goldstein, R.Z., Heitzeg, M., 2018. Adolescent brain cognitive development (ABCD) study: Overview of substance use assessment methods. Dev Cogn Neurosci 32, 80–96.

Liu, Z., Wang, Y., Gerig, G., Gouttard, S., Tao, R., Fletcher, T., Styner, M., 2010. Quality Control of Diffusion Weighted Images. Proc SPIE Int Soc Opt Eng 7628.

Logan, G.D., 1994. On the ability to inhibit thought and action: A users’ guide to the stop signal paradigm. Inhibitory processes in attention, memory, and language. Academic Press, San Diego, CA, US, pp. 189–239.

Luciana, M., Bjork, J.M., Nagel, B.J., Barch, D.M., Gonzalez, R., Nixon, S.J., Banich, M.T., 2018. Adolescent neurocognitive development and impacts of substance use: Overview of the adolescent brain cognitive development (ABCD) baseline neurocognition battery. Dev Cogn Neurosci 32, 67–79.

McDonald, C.R., Hagler, D.J., Jr., Girard, H.M., Pung, C., Ahmadi, M.E., Holland, D., Patel, R.H., Barba, D., Tecoma, E.S., Iragui, V.J., Halgren, E., Dale, A.M., 2010. Changes in fiber tract integrity and visual fields after anterior temporal lobectomy. Neurology 75, 1631–1638.

Mirzaalian, H., Ning, L., Savadjiev, P., Pasternak, O., Bouix, S., Michailovich, O., Karmacharya, S., Grant, G., Marx, C.E., Morey, R.A., Flashman, L.A., George, M.S., McAllister, T.W., Andaluz, N., Shutter, L., Coimbra, R., Zafonte, R.D., Coleman, M.J., Kubicki, M., Westin, C.F., Stein, M.B., Shenton, M.E., Rathi, Y., 2018. Multi-site harmonization of diffusion MRI data in a registration framework. Brain Imaging Behav 12, 284–295.

Moeller, S., Yacoub, E., Olman, C.A., Auerbach, E., Strupp, J., Harel, N., Ugurbil, K., 2010. Multiband multislice GE-EPI at 7 tesla, with 16-fold acceleration using partial parallel imaging with application to high spatial and temporal whole-brain fMRI. Magn Reson Med 63, 1144–1153.

Morgan, P.S., Bowtell, R.W., McIntyre, D.J., Worthington, B.S., 2004. Correction of spatial distortion in EPI due to inhomogeneous static magnetic fields using the reversed gradient method. J Magn Reson Imaging 19, 499–507.

Mueller, S.G., Weiner, M.W., Thal, L.J., Petersen, R.C., Jack, C., Jagust, W., Trojanowski, J.Q., Toga, A.W., Beckett, L., 2005. The Alzheimer’s disease neuroimaging initiative. Neuroimaging Clin N Am 15, 869–877, xi-xii.

Patton, G.C., Azzopardi, P., Kennedy, E., Coffey, C., Mokdad, A., 2017. Global Measures of Health Risks and Disease Burden in Adolescents. In: rd, Bundy, D.A.P., Silva, N., Horton, S., Jamison, D.T., Patton, G.C. (Eds.), Child and Adolescent Health and Development, Washington (DC).

Pierpaoli, C., Jezzard, P., Basser, P.J., Barnett, A., Di Chiro, G., 1996. Diffusion tensor MR imaging of the human brain. Radiology 201, 637–648.

Power, J.D., Barnes, K.A., Snyder, A.Z., Schlaggar, B.L., Petersen, S.E., 2012. Spurious but systematic correlations in functional connectivity MRI networks arise from subject motion. Neuroimage 59, 2142–2154.

Power, J.D., Mitra, A., Laumann, T.O., Snyder, A.Z., Schlaggar, B.L., Petersen, S.E., 2014. Methods to detect, characterize, and remove motion artifact in resting state fMRI. Neuroimage 84, 320–341.

Reuter, M., Fischl, B., 2011. Avoiding asymmetry-induced bias in longitudinal image processing. Neuroimage 57, 19–21.

Reuter, M., Rosas, H.D., Fischl, B., 2010. Highly accurate inverse consistent registration: a robust approach. Neuroimage 53, 1181–1196.

Reuter, M., Schmansky, N.J., Rosas, H.D., Fischl, B., 2012. Within-subject template estimation for unbiased longitudinal image analysis. Neuroimage 61, 1402–1418.

Reuter, M., Tisdall, M.D., Qureshi, A., Buckner, R.L., van der Kouwe, A.J., Fischl, B., 2015. Head motion during MRI acquisition reduces gray matter volume and thickness estimates. Neuroimage 107, 107–115.

Rimol, L.M., Panizzon, M.S., Fennema-Notestine, C., Eyler, L.T., Fischl, B., Franz, C.E., Hagler, D. J., Lyons, M.J., Neale, M.C., Pacheco, J., Perry, M.E., Schmitt, J.E., Grant, M.D., Seidman, L.J., Thermenos, H.W., Tsuang, M.T., Eisen, S.A., Kremen, W.S., Dale, A.M., 2010. Cortical thickness is influenced by regionally specific genetic factors. Biol Psychiatry 67, 493–499.

Rohde, G.K., Barnett, A.S., Basser, P.J., Marenco, S., Pierpaoli, C., 2004. Comprehensive approach for correction of motion and distortion in diffusion-weighted MRI. Magn Reson Med 51, 103–114.

Satterthwaite, T.D., Wolf, D.H., Loughead, J., Ruparel, K., Elliott, M.A., Hakonarson, H., Gur, R.C., Gur, R.E., 2012. Impact of in-scanner head motion on multiple measures of functional connectivity: relevance for studies of neurodevelopment in youth. Neuroimage 60, 623–632.

Segonne, F., Dale, A.M., Busa, E., Glessner, M., Salat, D., Hahn, H.K., Fischl, B., 2004. A hybrid approach to the skull stripping problem in MRI. Neuroimage 22, 1060–1075.

Segonne, F., Pacheco, J., Fischl, B., 2007. Geometrically accurate topology-correction of cortical surfaces using nonseparating loops. IEEE Trans Med Imaging 26, 518–529.

Seibert, T.M., Brewer, J.B., 2011. Default network correlations analyzed on native surfaces. J Neurosci Methods 198, 301–311.

Setsompop, K., Gagoski, B.A., Polimeni, J.R., Witzel, T., Wedeen, V.J., Wald, L.L., 2012. Blipped-controlled aliasing in parallel imaging for simultaneous multislice echo planar imaging with reduced g-factor penalty. Magn Reson Med 67, 1210–1224.

Siegel, J.S., Power, J.D., Dubis, J.W., Vogel, A.C., Church, J.A., Schlaggar, B.L., Petersen, S.E., 2014. Statistical improvements in functional magnetic resonance imaging analyses produced by censoring high-motion data points. Hum Brain Mapp 35, 1981–1996.

Sled, J.G., Zijdenbos, A.P., Evans, A.C., 1998. A nonparametric method for automatic correction of intensity nonuniformity in MRI data. IEEE Trans Med Imaging 17, 87–97.

Smith, S.M., Jenkinson, M., Woolrich, M.W., Beckmann, C.F., Behrens, T.E., Johansen-Berg, H., Bannister, P.R., De Luca, M., Drobnjak, I., Flitney, D.E., Niazy, R.K., Saunders, J., Vickers, J., Zhang, Y., De Stefano, N., Brady, J.M., Matthews, P.M., 2004. Advances in functional and structural MR image analysis and implementation as FSL. Neuroimage 23 Suppl 1, S208–219.

Thompson, W.K., Holland, D., 2011. Bias in tensor based morphometry Stat-ROI measures may result in unrealistic power estimates. Neuroimage 57, 1–4.

Tisdall, M.D., Hess, A.T., Reuter, M., Meintjes, E.M., Fischl, B., van der Kouwe, A.J., 2012. Volumetric navigators for prospective motion correction and selective reacquisition in neuroanatomical MRI. Magn Reson Med 68, 389–399.

Tisdall, M.D., Reuter, M., Qureshi, A., Buckner, R.L., Fischl, B., van der Kouwe, A.J.W., 2016. Prospective motion correction with volumetric navigators (vNavs) reduces the bias and variance in brain morphometry induced by subject motion. Neuroimage 127, 11–22.

Triantafyllou, C., Hoge, R.D., Krueger, G., Wiggins, C.J., Potthast, A., Wiggins, G.C., Wald, L.L., 2005. Comparison of physiological noise at 1.5 T, 3 T and 7 T and optimization of fMRI acquisition parameters. Neuroimage 26, 243–250.

Tukey, J.W., 1960. A survey of sampling from contaminated distributions. In: Olkin, I., Ghurye, S., Hoeffding, W., Madow, W., Mann, H. (Eds.), Contributions to Probability and Statistics. Stanford University Press, Stanford, pp. 448–485.

Uban, K.A., Horton, M.K., Jacobus, J., Heyser, C., Thompson, W.K., Tapert, S.F., Madden, P.A.F., Sowell, E.R., Adolescent Brain Cognitive Development, S., 2018. Biospecimens and the ABCD study: Rationale, methods of collection, measurement and early data. Dev Cogn Neurosci 32, 97–106.

Van Dijk, K.R., Hedden, T., Venkataraman, A., Evans, K.C., Lazar, S.W., Buckner, R.L., 2010. Intrinsic functional connectivity as a tool for human connectomics: theory, properties, and optimization. J Neurophysiol 103, 297–321.

Van Dijk, K.R., Sabuncu, M.R., Buckner, R.L., 2012. The influence of head motion on intrinsic functional connectivity MRI. Neuroimage 59, 431–438.

Van Essen, D.C., Ugurbil, K., Auerbach, E., Barch, D., Behrens, T.E., Bucholz, R., Chang, A., Chen, L., Corbetta, M., Curtiss, S.W., Della Penna, S., Feinberg, D., Glasser, M.F., Harel, N., Heath, A.C., Larson-Prior, L., Marcus, D., Michalareas, G., Moeller, S., Oostenveld, R., Petersen, S.E., Prior, F., Schlaggar, B.L., Smith, S.M., Snyder, A.Z., Xu, J., Yacoub, E., Consortium, W.U.-M.H., 2012. The Human Connectome Project: a data acquisition perspective. Neuroimage 62, 2222–2231.

Volkow, N.D., Koob, G.F., Croyle, R.T., Bianchi, D.W., Gordon, J.A., Koroshetz, W.J., Perez-Stable, E.J., Riley, W.T., Bloch, M.H., Conway, K., Deeds, B.G., Dowling, G.J., Grant, S., Howlett, K.D., Matochik, J.A., Morgan, G.D., Murray, M.M., Noronha, A., Spong, C.Y., Wargo, E. M., Warren, K.R., Weiss, S.R.B., 2018. The conception of the ABCD study: From substance use to a broad NIH collaboration. Dev Cogn Neurosci 32, 4–7.

Wald, L., Schmitt, F., Dale, A., 2001. Systematic spatial distortion in MRI due to gradient nonlinearities. Neuroimage 13, 50.

Wells, W.M., 3rd, Viola, P., Atsumi, H., Nakajima, S., Kikinis, R., 1996. Multi-modal volume registration by maximization of mutual information. Med Image Anal 1, 35–51.

Westlye, L.T., Walhovd, K.B., Dale, A.M., Espeseth, T., Reinvang, I., Raz, N., Agartz, I., Greve, D. N., Fischl, B., Fjell, A.M., 2009. Increased sensitivity to effects of normal aging and Alzheimer’s disease on cortical thickness by adjustment for local variability in gray/white contrast: a multi-sample MRI study. Neuroimage 47, 1545–1557.

White, N., Roddey, C., Shankaranarayanan, A., Han, E., Rettmann, D., Santos, J., Kuperman, J., Dale, A., 2010. PROMO: Real-time prospective motion correction in MRI using image-based tracking. Magn Reson Med 63, 91–105.

White, N.S., Leergaard, T.B., D’Arceuil, H., Bjaalie, J.G., Dale, A.M., 2013a. Probing tissue microstructure with restriction spectrum imaging: Histological and theoretical validation. Hum Brain Mapp 34, 327–346.

White, N.S., McDonald, C., Farid, N., Kuperman, J., Karow, D., Schenker-Ahmed, N.M., Bartsch, H., Rakow-Penner, R., Holland, D., Shabaik, A., Bjornerud, A., Hope, T., Hattangadi-Gluth, J., Liss, M., Parsons, J.K., Chen, C.C., Raman, S., Margolis, D., Reiter, R.E., Marks, L., Kesari, S., Mundt, A.J., Kane, C.J., Carter, B.S., Bradley, W.G., Dale, A.M., 2014. Diffusion-weighted imaging in cancer: physical foundations and applications of restriction spectrum imaging. Cancer Res 74, 4638–4652.

White, N.S., McDonald, C.R., Farid, N., Kuperman, J.M., Kesari, S., Dale, A.M., 2013b. Improved conspicuity and delineation of high-grade primary and metastatic brain tumors using “restriction spectrum imaging”: quantitative comparison with high B-value DWI and ADC. AJNR Am J Neuroradiol 34, 958–964, S951.

Yendiki, A., Koldewyn, K., Kakunoori, S., Kanwisher, N., Fischl, B., 2013. Spurious group differences due to head motion in a diffusion MRI study. Neuroimage 88C, 79–90.

Zaitsev, M., Maclaren, J., Herbst, M., 2015. Motion artifacts in MRI: A complex problem with many partial solutions. J Magn Reson Imaging 42, 887–901.

Zhuang, J., Hrabe, J., Kangarlu, A., Xu, D., Bansal, R., Branch, C.A., Peterson, B.S., 2006. Correction of eddy-current distortions in diffusion tensor images using the known directions and strengths of diffusion gradients. J Magn Reson Imaging 24, 1188–1193.

Zucker, R.A., Gonzalez, R., Feldstein Ewing, S.W., Paulus, M.P., Arroyo, J., Fuligni, A., Morris, A.S., Sanchez, M., Wills, T., 2018. Assessment of culture and environment in the Adolescent Brain and Cognitive Development Study: Rationale, description of measures, and early data. Dev Cogn Neurosci 32, 107–120.

